# STAT1–p53 Dynamics Program Cell Fate through p21 and PUMA

**DOI:** 10.1101/2025.10.28.685172

**Authors:** M.K. Francis Didier Tshianyi, Rebecca Walo Omana, Apollinaire Ndondo Mboma, Didier Kumwimba, Didier Gonze

**Affiliations:** Unit of Theoretical Chronobiology, Faculty of Sciences, Université libre de Bruxelles (ULB), Brussels, Belgium; Department of Mathematics and Computer Science, Faculty of Science and Technology, University of Kinshasa, Kinshasa, DR Congo; Department of Mathematics and Computer Science, Faculty of Sciences, University of Lubumbashi, Lubumbashi, DR Congo; Department of Science and Technologies, Faculty of Sciences and Technologies, New Horizon University, Lubumbashi, DR Congo

**Keywords:** Hopf-bifurcation, Oscillations, Cell-fate, Stress, p53-Mdm2, STAT1, Senescence, Apoptosis

## Abstract

Signalling pathways tightly regulate stress-induced cell fate decisions, with *p*53 stabilization via attenuation of the *p*53–*Mdm*2 feedback being central to effective responses. Signal Transducer and Activator of Transcription 1 (STAT1) both represses *Mdm2* transcription and co-activates p53 targets (*p21* and *PUMA*), yet these dual roles have not been quantified within a unified framework. We build a mechanistic model that couples a biologically calibrated stress scale (*S*) and STAT1 activity (Σ) to the canonical *p*53–*Mdm*2 core and to downstream *p21* and *PUMA* modules. Key processes are not described by fixed parameters but by explicit functions of *S*, allowing the model to self-adjust across stress levels without manual re-fitting. This formulation ensures that oscillatory, damped, or plateau responses naturally emerge by varying *S* alone. STAT1 acts as a dynamical gain on the core (controlling mean level, pulse amplitude, and duration) and as a transcriptional co-activator scaling promoter strengths. Bifurcation analysis reveals a two-dimensional Hopf region in the (*S*, Σ) plane; increasing Σ shifts this region to higher *S* and progressively narrows it, ultimately quenching oscillations at large Σ . Simulations of a *generic* cell with time-varying *S* and Σ, combined with a two-stage decision rule (early transient detection followed by stationary readout), map *p*53 dynamics to fate: sustained moderate oscillations align with arrest, damped intermediate responses with senescence, and strongly damped high plateaus with apoptosis. The model reproduces cell fate distributions reported in literature for different cell lines (MCF-7, HCT116, U2OS) without kinetic parameter re-fitting, and highlights cell-type-specific sensitivity to *p21* versus *PUMA*. Our framework identifies STAT1 as a tunable amplifier and oscillation quencher of the stress-responsive p53 network, providing testable dynamics-based predictions for fate control.

## Introduction

Cellular stress can profoundly reshape protein dynamics. It arises from diverse agents (UV, *γ*-irradiation, viral infection…) and perturbs genome integrity and homeostasis. To cope with these adverse conditions, cells deploy signalling networks in which the tumour suppressor *p53* is central. Under basal conditions, *p*53 is kept low by its principal regulator *Mdm2*, which binds and ubiquitinates *p*53, promoting nuclear export or proteasomal degradation; this forms a negative feedback in which *p*53 induces *Mdm2*, and *Mdm2* accelerates *p*53 turnover (Bar-Or et al., 2000; Lahav et al., 2004; Ciliberto et al., 2005; Proctor & Gray, 2008). A hallmark of this module is its ability to generate *oscillations* in response to DNA damage, observed experimentally and captured by prior models (Abou-Jaoudé et al., 2009; Ouattara et al., 2010; Batchelor et al., 2011).

Beyond the canonical DNA-damage axis, we consider the transcription factor STAT1. Biologically, STAT1 is activated by cytokines (notably type I/II interferons) via JAK/STAT signalling; upon activation it dimerizes, translocates to the nucleus and modulates gene expression (Levy & Darnell, 2002; Giles et al., 2017). Importantly, studies indicate that STAT1 can stabilise *p*53 and cooperate to regulate the expression of *p*53 target genes (Brzostek-Racine et al., 2011; Yu et al., 2015; Heitmeier et al., 1999). In practical terms, this gives two conceptually distinct inputs: a *stress/DNA-damage* signal (e.g. ATM/Chk-mediated) and a *cytokine/STAT1* signal (e.g. interferon bursts). These inputs can combine in time—for instance, after irradiation the effective DNA damage may *decrease* as lesions are repaired while paracrine interferon induces a *transient STAT1 peak*; conversely, inflammatory contexts can sustain STAT1 activity with minimal ongoing damage.

It is not straightforward to predict how different mixtures of stress and STAT1 activity, acting through *p*53, will shape the dynamics of downstream protein effectors, namely *p21* and *PUMA*, nor how cells interpret these signals to choose between *cell-cycle arrest, senescence* or *apoptosis*. Although biology suggests that STAT1 represses *Mdm2* and enhances *p*53 target programmes (Brzostek-Racine et al., 2011; Feng et al., 2020);, to our knowledge, no mathematical studies have explicitly integrated STAT1 with the *p*53–*Mdm*2 oscillator and its downstream effectors. This gap underlies the need for a kinetic model.

We develop a mechanistic model of the canonical *p*53–*Mdm*2 core that explicitly takes the stress level *S* and STAT1 activity Σ as inputs and propagates their effects to the two key effectors *p21* and *PUMA*. In the model, STAT1 both *represses Mdm2* transcription (stabilising *p*53) and *co-activates p*53 targets (*p21, PUMA*). We introduce a two-stage fate criterion: (i) an *early* transient detector marking commitment once *p21* /*PUMA* exceed thresholds for a minimal dwell time, and (ii) a *late* stationary assessment within 60–72 h using the same threshold-based rules. A bifurcation analysis reveals a two-dimensional oscillatory region delimited by Hopf bifurcations in the (*S*, Σ) plane that organises transitions between oscillatory and plateau regimes of *p*53. In a single “standard” cell under time-constant *S*, increasing Σ precipitates the early detector and shortens decision times, biasing outcomes toward the *PUMA*-dominated (pro-apoptotic) branch; low–moderate Σ favours *p21* -driven arrest/senescence. At the population level, we apply the same rule—without re-fitting biochemical parameters—to archetypes cell lines (MCF-7, HCT116, U2OS), obtaining literature-consistent fate splits and revealing cell-type-specific sensitivity to *p21* vs *PUMA* (Abbas & Dutta, 2009; Jeffers et al., 2003). Throughout, qualitative *p*53 motifs map to fate tendencies (sustained moderate-mean oscillations are linked to cell cycle arrest; damped intermediate responses are linked to senescence; strongly damped high plateaux are linked to apoptosis), providing testable predictions under time-varying *S* and Σ.

Most mathematical models of the *p*53–*Mdm*2 module assume *fixed* parameters. To reproduce different behaviours under low, moderate, or high stress, one typically has to re-tune rate constants or re-fit parameter sets for each condition (see e.g. (Bar-Or et al., 2000; Proctor & Gray, 2008; Ciliberto et al., 2005; Eliaš et al., 2014)). By contrast, our framework is intrinsically *stress-modulated*: the input *S* explicitly shapes several key processes through functional dependencies (Hill-type or saturating terms). As a consequence, no additional manual adjustment is required to generate the repertoire of dynamical responses. Our model allows to predict cell fate under various profiles of stress and Stat1.

We integrate a biologically calibrated stress scale with a dual-role STAT1 modulation of the *p*53–*Mdm*2 network, introduce a decision rule that scales from single cells to heterogeneous populations, and reveal a two-dimensional Hopf region in (*S*, Σ) that highlights oscillation–plateau transitions. Our main contributions are:

i. Integrating the dual-role STAT1 in one mechanistic framework: repression of *Mdm2* (stabilizing *p*53) and co-activation of *p21* /*PUMA*.
ii. Identifying the two-dimensional Hopf region in (*S*, Σ)that shifts rightward and narrows with Σ, reshaping oscillatory domains of *p*53.
iii. Implementation of Stress-modulated parameters that embed *S* directly into the system’s kinetics, eliminating the need for manual re-tuning across stress levels.
iv. Defining a two-stage fate rule (early detector + late stationary readout) mapping *p*53 dynamics to arrest/senescence/apoptosis, with thresholds and minimal dwell times.
v. A population-level validation across archetypes (MCF-7, HCT116, U2OS) *without biochemical re-fitting*, reproducing literature distributions and revealing sensitivity to *p21* versus *PUMA*.

Section 1 introduces the model and the assumptions used; Section 2 presents the bifurcation analysis and *p*53 dynamical patterns; Section 3 formalises the two-stage fate rule and illustrates their application on single-cells; Section 4.1 introduces the archetypes, their calibration targets, and demonstrates the two-stage fate rule on them. Section 4.4 discusses biological implications, limitations, and testable predictions.

## 1. The model

In this section we present the hypothesis and the different processes we used to construct our model.

We consider a minimal *p*53–*Mdm2* module driven by two *external inputs*: the DNA-damage/stress signal *S*(*t*) and the STAT1 activity Σ(*t*). In this module they are treated as prescribed forcings (constants or time profiles), i.e. as parameters rather than state variables.

*p53 balance. p*53 is produced at a basal rate *β*_0_ and is further induced by stress through a sigmoidal Hill term of order 4, 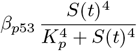. Removal of *p*53 by *Mdm2* is described by a Michaelis–Menten term 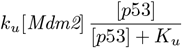, multiplied by 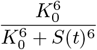 so that increasing *S weakens* the effective ubiquitylation capacity and stabilises *p*53. A first–order term *δ*_*p*_[*p*53] accounts for basal turnover.

*Mdm2 RNA and protein*. Transcription of *Mdm2* mRNA is activated by *p*53 with *apparent cooperativity* (quadratic dependence) and *repressed* by STAT1,

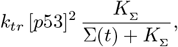

where the quadratic [*p*53]^2^ term follows (Hunziker et al., 2010), who used it to capture the effective cooperativity/tetramerisation of *p*53 in driving *Mdm2* expression. The mRNA decays at rate *δ*_*m*_. *Mdm2* protein is translated at rate *k*_*tl*_[*Mdm2*_*RNA*_] and degraded with rate *δ*_*M*_ [*Mdm2*].

The model reads:

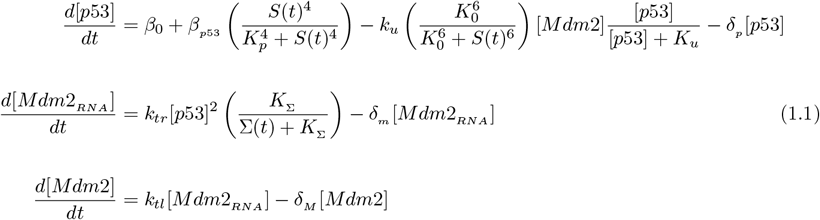

Throughout the manuscript, square brackets denote *concentrations*. For any molecular species *X*, we write [*X*] for its concentration (typically in nM or *µ*M), e.g. [*p*21], [*PUMA*], [*p*53]. When time dependence is relevant we use [*X*](*t*). For readability, the ODE state variables may be written without brackets (e.g., *p*21) but are understood to represent the same concentration quantity with the same units unless explicitly stated otherwise.

### 1.1 Stress scale

Stress (*S*) is a single, dimensionless control parameter that modulates all stress–sensitive processes in the model, including the *p*53 half-life via ubiquitination, as well as the production of *p*53, *p*21, and PUMA. To enable comparison across experiments, we calibrated a simple scale *S* ∈ [0, 25] from published responses to UV and *γ*-irradiation (1). These results were calibrated primarily based on:

(i) non-transformed murine cells (UV); (ii) human skin *in vivo* (UV); and (iii) the human cell lines MCF-7, MCF10A, and RPE-hTERT for fine measurements of dynamics and half-life (UV and *γ*). In the manuscript, *S* = 0 denotes no insult and larger values encode stronger damage.

We will use the term *generic* cell or *standard* cell to refer to any cell that reproduces the behaviour described in Table 1.

**Table 1.** Calibrated stress scale *S*: correspondence with benchmark *γ*-irradiation doses, qualitative biological effects, and *p*53 dynamics. UV exposures were mapped to *S* by matching *p*53 stabilization metrics (see text).

| $S$ | Benchmark Gy | Main Biological Effect | $p53$ Dynamics | References |
| --- | --- | --- | --- | --- |
| 0 | 0 | No activation, efficient repair | Very low, stable | Bar-Or et al. (2000); Lahav et al. (2004) |
| 2–5 | 1.5–4 | Repairable damage; minimal p21 activation | Transient, low amplitude | Batchelor et al. (2008); Geva-Zatorsky et al. (2006) |
| 6.5 | 5 | Cell-cycle arrest; clear p21 activation | Moderate, damped oscillations | Geva-Zatorsky et al. (2006); Purvis et al. (2012) |
| 10 | 8 | Mixed p21/PUMA activation, bifurcated response | Robust oscillations | Lahav et al. (2004); Stewart-Ornstein & Lahav (2017) |
| 14 | 11 | Likely apoptosis via PUMA/BAX | Large then damped oscillations | Nakano & Vousden (2001); Jeffers et al. (2003) |
| 18 | 14 | Full apoptosis or terminal senescence | High stabilization, end of oscillations | Purvis et al. (2012); Stewart-Ornstein & Lahav (2017) |
| 21 | 17 | Massive apoptosis, irreversible stress | High plateau, lost dynamics | Jeffers et al. (2003) |
| 25 | 20+ | Loss of viability, collapse | Unstable/absent $p53$ | Purvis et al. (2012) |

**Table 2.**
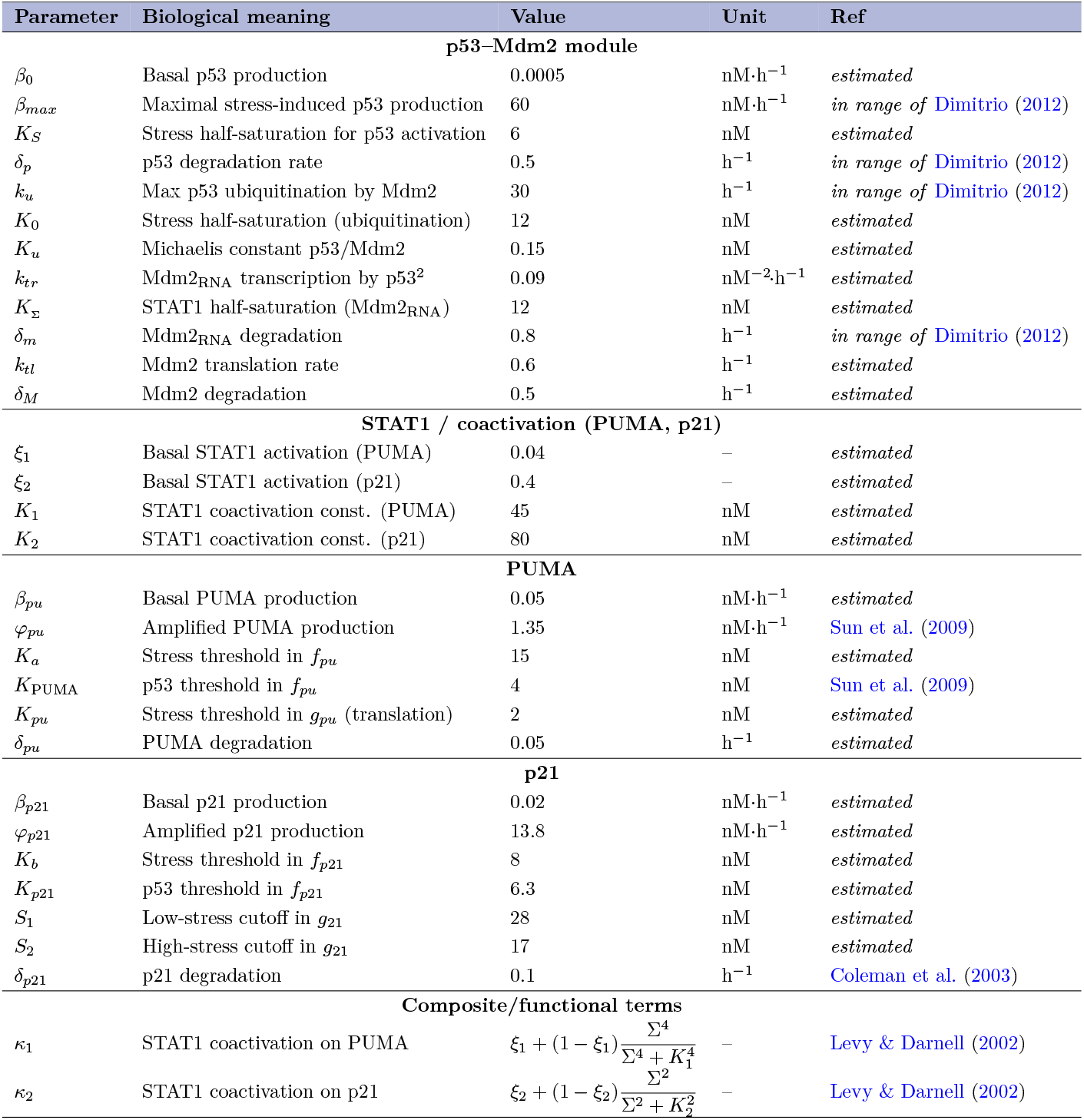
Parameters for the p53–Mdm2–p21–PUMA model. In **Ref**, *estimated* indicates no published numeric interval was found that contains the table value. The parameter values given here account for a *generic* cell.

| Parameter | Biological meaning | Value | Unit | Ref |
| --- | --- | --- | --- | --- |
| **p53–Mdm2 module** |  |  |  |  |
| $\beta_0$ | Basal p53 production | 0.0005 | nM·h<sup>-1</sup> | *estimated* |
| $\beta_{max}$ | Maximal stress-induced p53 production | 60 | nM·h<sup>-1</sup> | *in range of* Dimitrio (2012) |
| $K_S$ | Stress half-saturation for p53 activation | 6 | nM | *estimated* |
| $\delta_p$ | p53 degradation rate | 0.5 | h<sup>-1</sup> | *in range of* Dimitrio (2012) |
| $k_u$ | Max p53 ubiquitination by Mdm2 | 30 | h<sup>-1</sup> | *in range of* Dimitrio (2012) |
| $K_0$ | Stress half-saturation (ubiquitination) | 12 | nM | *estimated* |
| $K_u$ | Michaelis constant p53/Mdm2 | 0.15 | nM | *estimated* |
| $k_{tr}$ | Mdm2<sub>RNA</sub> transcription by p53<sup>2</sup> | 0.09 | nM<sup>-2</sup>·h<sup>-1</sup> | *estimated* |
| $K_\Sigma$ | STAT1 half-saturation (Mdm2<sub>RNA</sub>) | 12 | nM | *estimated* |
| $\delta_m$ | Mdm2<sub>RNA</sub> degradation | 0.8 | h<sup>-1</sup> | *in range of* Dimitrio (2012) |
| $k_{tl}$ | Mdm2 translation rate | 0.6 | h<sup>-1</sup> | *estimated* |
| $\delta_M$ | Mdm2 degradation | 0.5 | h<sup>-1</sup> | *estimated* |
| **STAT1 / coactivation (PUMA, p21)** |  |  |  |  |
| $\xi_1$ | Basal STAT1 activation (PUMA) | 0.04 | – | *estimated* |
| $\xi_2$ | Basal STAT1 activation (p21) | 0.4 | – | *estimated* |
| $K_1$ | STAT1 coactivation const. (PUMA) | 45 | nM | *estimated* |
| $K_2$ | STAT1 coactivation const. (p21) | 80 | nM | *estimated* |
| **PUMA** |  |  |  |  |
| $\beta_{pu}$ | Basal PUMA production | 0.05 | nM·h<sup>-1</sup> | *estimated* |
| $\varphi_{pu}$ | Amplified PUMA production | 1.35 | nM·h<sup>-1</sup> | Sun et al. (2009) |
| $K_a$ | Stress threshold in $f_{pu}$ | 15 | nM | *estimated* |
| $K_{PUMA}$ | p53 threshold in $f_{pu}$ | 4 | nM | Sun et al. (2009) |
| $K_{pu}$ | Stress threshold in $g_{pu}$ (translation) | 2 | nM | *estimated* |
| $\delta_{pu}$ | PUMA degradation | 0.05 | h<sup>-1</sup> | *estimated* |
| **p21** |  |  |  |  |
| $\beta_{p21}$ | Basal p21 production | 0.02 | nM·h<sup>-1</sup> | *estimated* |
| $\varphi_{p21}$ | Amplified p21 production | 13.8 | nM·h<sup>-1</sup> | *estimated* |
| $K_b$ | Stress threshold in $f_{p21}$ | 8 | nM | *estimated* |
| $K_{p21}$ | p53 threshold in $f_{p21}$ | 6.3 | nM | *estimated* |
| $S_1$ | Low-stress cutoff in $g_{21}$ | 28 | nM | *estimated* |
| $S_2$ | High-stress cutoff in $g_{21}$ | 17 | nM | *estimated* |
| $\delta_{p21}$ | p21 degradation | 0.1 | h<sup>-1</sup> | Coleman et al. (2003) |
| **Composite/functional terms** |  |  |  |  |
| $\kappa_1$ | STAT1 coactivation on PUMA | $\xi_1 + (1 - \xi_1) \frac{\Sigma^4}{\Sigma^4 + K_1^4}$ | – | Levy & Darnell (2002) |
| $\kappa_2$ | STAT1 coactivation on p21 | $\xi_2 + (1 - \xi_2) \frac{\Sigma^2}{\Sigma^2 + K_2^2}$ | – | Levy & Darnell (2002) |

### 1.2 Stat1 and stress

In this article Σ(*t*) denotes an interferon (IFN)–driven JAK–STAT signal that activates STAT1 (Y701 phosphorylation, dimerization, nuclear translocation). Σ(*t*) captures the *cytokine context* (paracrine/autocrine IFN-*α/β* or IFN-*γ*) that co-activates parts of the *p*53 program, including PUMA and *p*21 (Levy & Darnell, 2002; Giles et al., 2017; Kalliara et al., 2022; Yu et al., 2015; Abbas & Dutta, 2009; Nakano & Vousden, 2001).

A single insult (irradiation, UV, some chemotherapies) can *simultaneously* activate the *p*53 arm via DNA damage and, through transient interferon bursts, the JAK–STAT pathway and STAT1; in such coupled cases, *S* and Σ rise together but with distinct kinetics (often a delayed Σ) (Bar-Or et al., 2000; Lahav et al., 2004; Proctor & Gray, 2008; Batchelor et al., 2011). Conversely, cytokine exposure or infection can elevate Σ with little DNA damage, while sterile genotoxic stress can raise *S* with minimal interferon signalling (Vousden & Ryan, 2009). These scenarios motivate treating *S*(*t*) and Σ(*t*) as *independent, externally controlled inputs* and exploring their combinations/timings to reveal how they reshape *p*53 dynamics and the downstream balance between *p21* and PUMA.

### 1.3 p21 and PUMA

*p21* (*CDKN1A*; p21^Cip1*/*Waf1^) is the cyclin–dependent kinase inhibitor *CDKN1A*, historically referred to as *p*21^Cip1*/*Waf1^ (also SDI1). It promotes cell–cycle arrest and, under sustained activation, senescence. In the *p*53 network, *p*21 acts as a relatively *slow*, integrative branch: its mRNA/protein turnover is slower than immediate–early responders, so its level reflects the time–integrated history of *p*53 activity (El-Deiry et al., 1993; Harper et al., 1993; Xiong et al., 1993; Abbas & Dutta, 2009; Barr et al., 2017; Jin et al., 2003; Coleman et al., 2003) .

*PUMA* stands for *p53 Upregulated Modulator of Apoptosis*. It is a *p*53–induced BH3–only protein that antagonizes anti–apoptotic BCL–2 family members. Functionally, PUMA is a *fast*, switch–like effector: it can rise quickly following strong *p*53 activity and directly commits cells to apoptosis when its threshold is crossed (Nakano & Vousden, 2001; Jeffers et al., 2003; Purvis et al., 2012; Hanson et al., 2019; Yu et al., 2015) . We extend the core *p*53–*Mdm*2 module in (1.1) with a branch for *p*21 and PUMA.

*p21 (cell-cycle arrest/senescence)*. To capture *p*21’s early engagement under mild damage, stress *S*(*t*) enhances transcription through a low-threshold sigmoidal term,

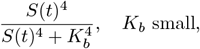

so that half-activation occurs at small *S* (Abbas & Dutta, 2009; Stewart-Ornstein & Lahav, 2017; Barr et al., 2017). At stronger stress, multiple post-transcriptional/post-translational checks are known to limit *p*21 (e.g., altered stability and proteasomal turnover) (Jin et al., 2003; Coleman et al., 2003; Hanson et al., 2019; Schwanhäusser et al., 2011). We therefore include a stress-dependent translational factor

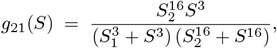

which is near-maximal at intermediate *S* but *attenuates p*21 production at very high *S*, reflecting the reported dampening of *p*21 under strong stimuli. Regarding STAT1, we allow only a *modest* co-activation of *p*21, consistent with literature indicating that *p*21 is primarily governed by *p*53 at low–moderate stress while STATs can provide auxiliary input (Levy & Darnell, 2002; Abbas & Dutta, 2009; Yu et al., 2015).

*PUMA (apoptosis)*. PUMA is a pro-apoptotic *p*53 target that is robustly induced under genotoxic stress (Nakano & Vousden, 2001; Jeffers et al., 2003). In contrast to *p*21, we posit a higher transcriptional threshold for stress,

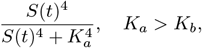

so that PUMA half-activation occurs at stronger *S*, aligning with its role in apoptosis (Purvis et al., 2012; Stewart-Ornstein & Lahav, 2017). To model stress resilience of translation, we include a gain

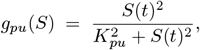

motivated by stress-sensitive translational control observed in the *p*53 network and proteome-wide measurements (Schwanhäusser et al., 2011; Hanson et al., 2019). Unlike *p*21, PUMA is assumed to be *more sensitive* to STAT1: we multiply its promoter gain by a steeper co-activation factor of Σ (e.g. a higher Hill sensitivity), consistent with interferon/STAT1 signalling intersecting pro-apoptotic *p*53 programmes (Levy & Darnell, 2002; Giles et al., 2017; Brzostek-Racine et al., 2011). The extended model is given by:

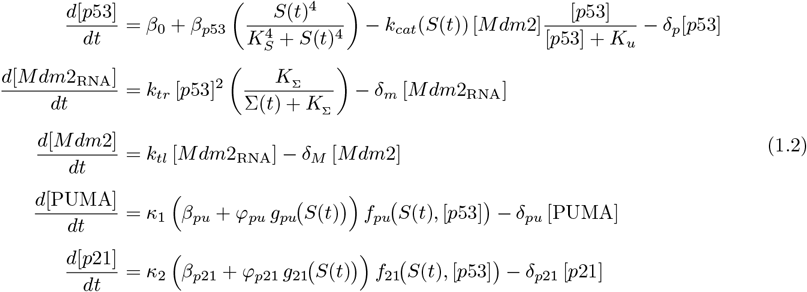

Where:

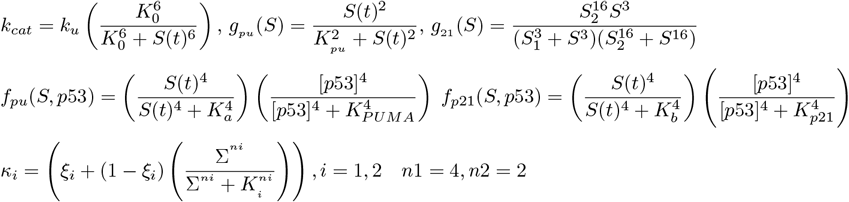

The following table gives the parameters values and their references:

## 2. Stability and bifurcation

In this section, we characterise both the *dynamical* behaviour and the *mathematical* properties of the model that couples stress (*S*) and STAT1 activity (Σ) to the canonical *p*53–*Mdm*2 core and its downstream effectors *p21* and *PUMA*.

### 2.1. Stress–dependent dynamics of the p53–Mdm2 core

We first examined how the core module responds when only the *stress* input *S* is varied (at fixed STAT1 activity Σ = 100). Figure 2 illustrates the time courses of *p*53, *Mdm*2_*RNA*_ and *Mdm*2 protein for representative levels *S* ∈ {2, 10, 18, 25} . All simulations are initialized close to the low-stress steady state of the model. Specifically, each species starts from a small but non-zero basal concentration (*p*53 = 10^*−*3^, *Mdm*2_*RNA* = 10^*−*7^, *Mdm*2 = 1.5 × 10^*−*7^, *PUMA* = 10^*−*6^, *p*21 = 10^*−*6^). These values approximate the quasi-equilibrium reached in the absence of stress (*S* ≈ 0), where *p*53 is maintained at a low level by efficient Mdm2-mediated degradation, and both *PUMA* and *p21* remain transcriptionally silent. They prevent numerical artifacts associated with strictly zero initial conditions while preserving the biological interpretation of a basal, unstressed cell. Unless otherwise stated, these initial conditions are used for all single-cell simulations and results presented in this work.

**Figure 1.**
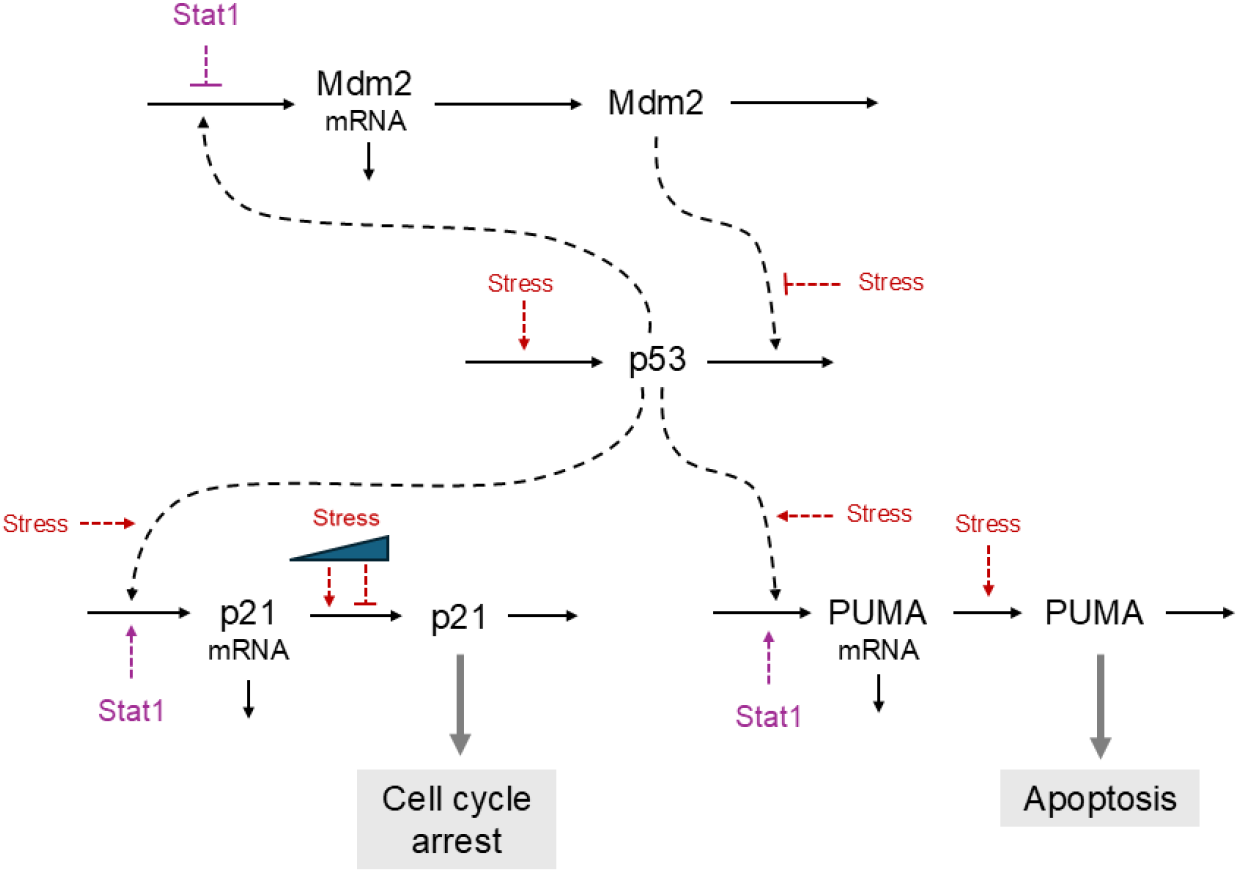
Scheme of the model: Red dashed labels (“Stress”) denote the stress input *S*(*t*), which stabilises *p*53 and modulates downstream processes; purple dashed labels (“Stat1”) denote interferon/STAT1 activity Σ(*t*). *p*53 activates *Mdm2* mRNA transcription, and *Mdm2* protein induces *p*53 degradation. STAT1 has a dual role: it represses *Mdm2* transcription (bar placed on the *p*53 → *Mdm2* mRNA link) and co-activates the *p*53 programme toward *p21* (cell-cycle arrest) and *PUMA* (apoptosis). Arrows indicate activation; blunt arrows indicate inhibition; dashed connections denote regulatory influences (not material flow). The triangular wedge on the p21 mRNA → *p*21 arrow encodes a stress-dependent translation gate: at low *S*, the effective translation rate decreases as *S* increases, and it shuts off for extreme stress levels.

**Figure 2.**
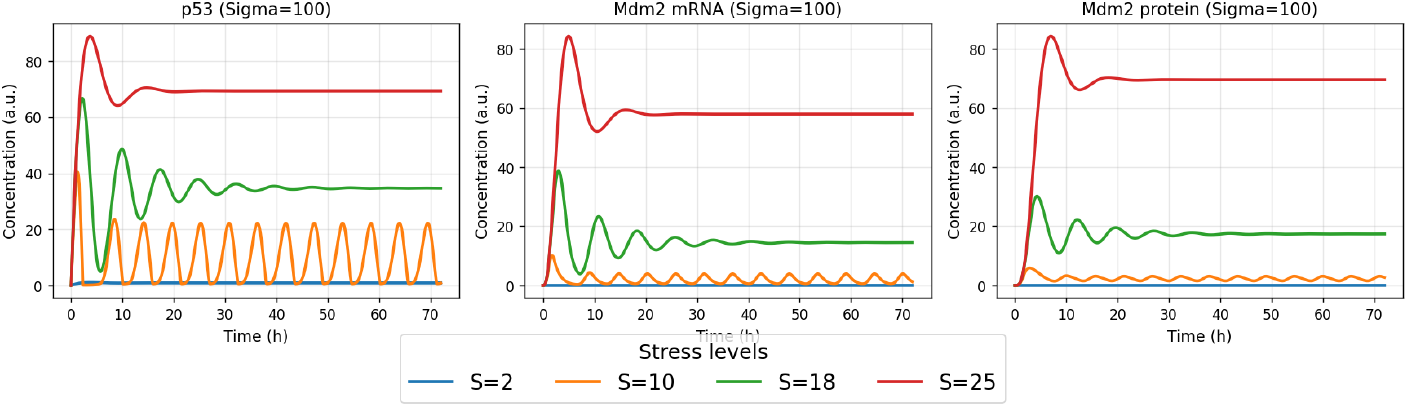
Stress scan at fixed Σ = 100. Time courses of *p*53, *Mdm*2_*RNA*_ and *M dm*2 protein for *S* ∈ {2, 10, 18, 25} . Low *S*: basal fixed point; intermediate *S*: sustained oscillations (period ∼ 6–8 h); high *S*: damped transients to an elevated plateau.

#### We have three dynamical regimes

For Low stress (e.g. *S* ≲ 4; *S* = 2 shown). The system remains close to basal: *p*53 is quickly cleared by *Mdm*2 and trajectories converge to a low steady state with negligible oscillations.

At Intermediate stress (oscillatory band; *S* = 10). We observe sustained *p*53 oscillations, mirrored by delayed oscillations of *Mdm*2 *RNA* and protein. The mean *p*53 level rises with *S*, the peak–to–peak amplitude is maximal in this band, and the inter-peak interval (IPI) is about 6.1 h. At High stress (damped/plateau; *S* = 18–25). A large overshoot is followed by damped oscillations that settle to an elevated plateau; for very high stress (*S* = 25), the response is essentially monotone. Here, the rising Mdm2 synthesis rapidly restores control, suppressing sustained oscillations.

#### Effect of S on p53 period, amplitude, and peak count

The global sensitivity (*SRC*, standardized regression coefficient, final–mean), presented in Section S2.2 of the Supplement, quantifies the normalized linear influence of each parameter on a given output variable across simulations. The SRC analysis is consistent with these trends: the period increases with *S* (positive SRC), the peak count decreases (negative SRC), and the stationary amplitude shows a weaker, overall negative dependence on *S*—in line with the large initial transient followed by diminished sustained oscillations at high stress. A bifurcation analysis will pinpoint the Hopf threshold *S*^*\**^ that separates the pulsatile regime from the damped, high-plateau regime.

#### Mechanistic interpretation

Stress acts on two levers of the core loop: (i) it *increases p*53 production through the sigmoidal term 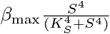; and (ii) it *weakens Mdm*2-mediated removal via the factor 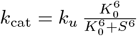. The core module exhibits behaviours like the one summarized in Table 1.

### 2.2. Existence, positivity, and boundedness of solutions

We consider the full five–dimensional system

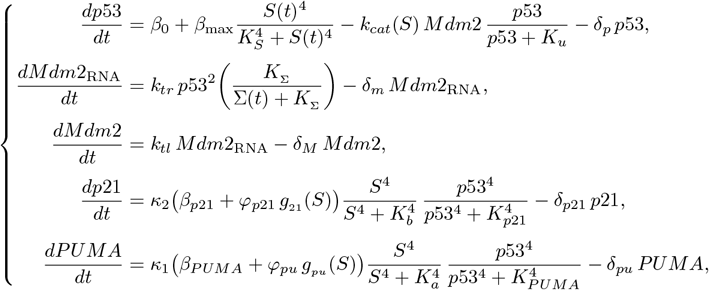

defined on 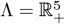, with strictly positive parameters and initial conditions. Here *g*_21_ and *g*_*pu*_ denote bounded Hill-type functions satisfying: 0 ≤ *g*_21_, *g*_*pu*_ ≤ 1.

**Theorem: Positivity and boundedness of the full model Theorem: Positivityl model**

For any nonnegative measurable inputs *S*(*t*), *Stat*1(*t*) and strictly positive initial conditions, the system above admits a unique global solution (*p*53, *Mdm*2_RNA_, *Mdm*2, *p*21, *PUMA*)(*t*) such that:

- The positive cone 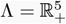 is invariant (all variables remain nonnegative).
- The trajectories are uniformly bounded in time: there exist explicit constants *C*_*p*53_, *C*_*mRNA*_, *C*_*Mdm*2_, *C*_*p*21_, *C*_*PUMA*_ depending only on the parameters such that *p*53(*t*) ≤ *C*_*p*53_, *Mdm*2_RNA_(*t*) ≤ *C*_*mRNA*_, *Mdm*2(*t*) ≤ *C*_*Mdm*2_, *p*21(*t*) ≤ *C*_*p*21_, *PUMA*(*t*) ≤ *C*_*PUMA*_, ∀*t* ≥ 0.

#### Sketch of proof. Positivity

At each boundary face (e.g. *p*53 = 0, *Mdm*2_RNA_ = 0, *p*21 = 0, *PUMA* = 0), the corresponding right-hand side is nonnegative, so that the positive cone Λ is invariant.

#### Bound for p53 (by the Grönwall–Bellman inequality)

Since 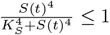 and 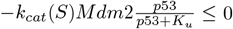, we obtain

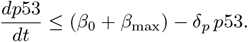

By the differential Grönwall inequality,

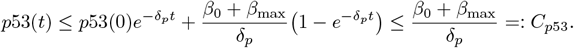

#### Bound for p21 (Grönwall)

The Hill terms satisfy 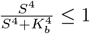 and 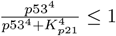, while *g*_21_ (*S*) ≤ 1.

Hence

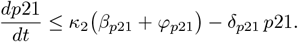

By Grönwall,

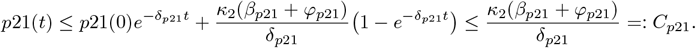

#### Other variables

For *Mdm*2_RNA_, *Mdm*2 and *PUMA*, one uses the previously obtained bound on *p*53 together with the fact that the Hill functions are bounded by 1. Each equation is of the form *u*^*′*^(*t*) ≤ *A* − *Bu*(*t*) with *A, B >* 0, so the Grönwall–Bellman inequality yields uniform bounds (full details and explicit constants *C*_*mRNA*_, *C*_*Mdm*2_, *C*_*PUMA*_ are given in the Supplement). Moreover, since the right-hand side is 𝒞 ^1^ in the state variables on Λ, the Picard–Lindelöf theorem ensures existence and *uniqueness* of a local solution. This establishes that the problem is *well posed* both mathematically and biologically.

#### Conclusion

The entire five-dimensional model is well-posed: all species concentrations remain nonnegative and bounded for all time. The Grönwall–Bellman inequality’s provides explicit bounds for *p*53 and *p*21; the other cases are analogous and deferred to the Supplement for clarity.

### 2.3. Stability

#### No bistability

For physiological consistency and a history–independent response, we must ensure that the system does not admit bistability. This is guaranteed by the following results:

*Equilibria (characterization)*.. Any equilibrium (*p*53^*\**^, *Mdm*2^*\**RNA^, *Mdm*2^*\**^, *p*21^*\**^, *PUMA*^*\**^) satisfies

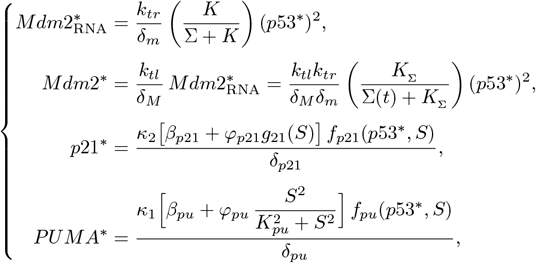

with *p*53^*\**^ solving

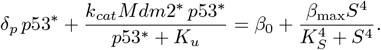

**Theorem: Uniqueness of the equilibrium point**

Consider the subsystem

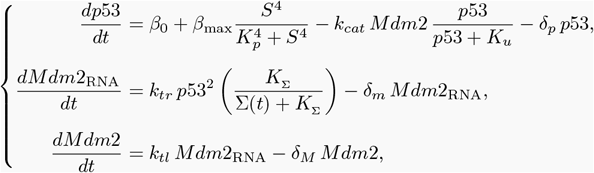

defined on 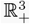 with strictly positive parameters and initial data.

Then, for any *S >* 0, the system admits a **unique** strictly positive equilibrium, obtained as the unique solution of

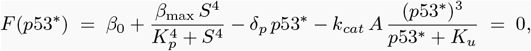

where 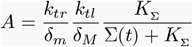 (a positive constant), and *F* is strictly decreasing.

Hence, there is neither bistability nor multiplicity of equilibria for any *S*.

#### Sketch of proof

At equilibrium one has 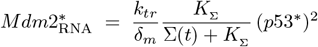 and 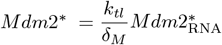, so the *p*53 steady-state equation reduces to *F* (*p*53^*\**^) = 0 with

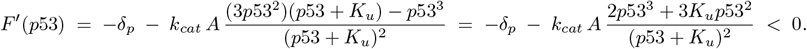

Thus *F* is strictly decreasing on (0, ∞). Since 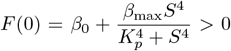 and *F* (*p*53) → −∞ as *p*53 → ∞, the intermediate value theorem yields a *unique* root *p*53^*\**^ *>* 0. Back–substitution gives a unique 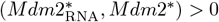. □

*Why this suffices for the full model*. In the full five-dimensional model, every equilibrium takes the form

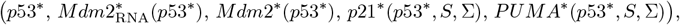

i.e., *all components are functions of p*53^*\**^. In particular, there can be no equilibrium multiplicity and hence no classical bistability of equilibria.

#### 3D phase portrait and stability

We fix STAT1 at Σ = 100 and track the Jacobian spectrum at the global equilibrium as stress *S* varies. The real part of the dominant complex pair crosses zero at *S*^*\**^ ≈ 4.32, indicating a supercritical Hopf bifurcation (Fig. S2). For *S < S*^*\**^, all species converge to a stable equilibrium; for *S > S*^*\**^, a stable limit cycle emerges. At Σ = 100 (Fig. 3), each column corresponds to a fixed value of *S*, while the rows display (Mdm2_RNA, Mdm2, *p*53), (Mdm2, *p*53, *p*21), and (Mdm2, *p*53, PUMA). Tight inward spirals indicate convergence toward a stable fixed point, whereas tubular coils correspond to stable limit cycles characterized by the expected phase lags, with *p*53 leading Mdm2 and *p*21/PUMA responding with additional delay.

**Figure 3.**
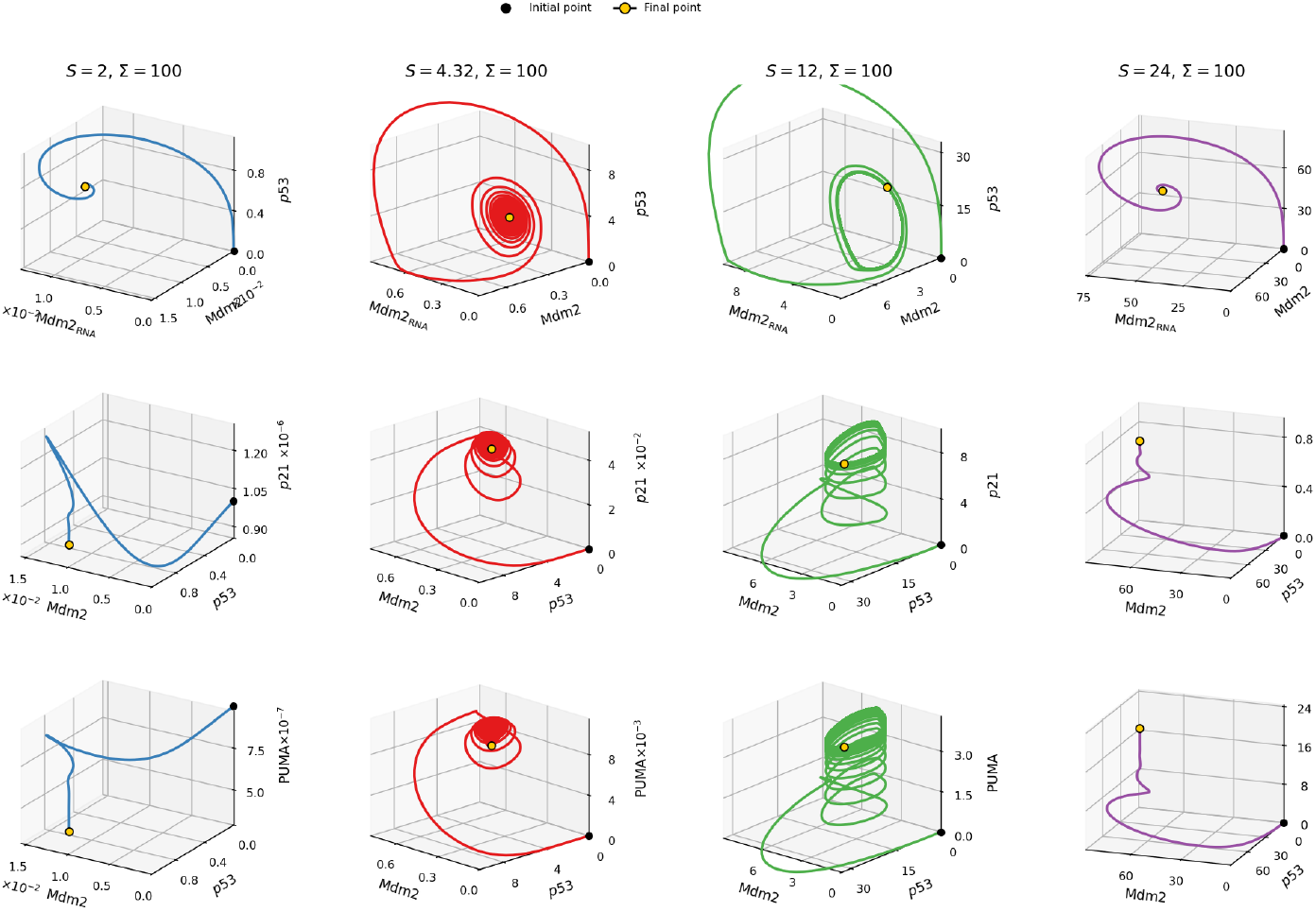
3D phase portraits for *p*21 and PUMA at *S* = 2, 4.32, 12, 24 with Σ = 100.

At *S* = 2 (low stress), the trajectories rapidly relax to a low fixed point: *p*53 is efficiently cleared by Mdm2 (reflecting high *k*_cat_ values), and both *p*21 and PUMA remain far below their activation thresholds. This regime corresponds to a cycling, unstressed state.

For *S* = 4.32 (near the Hopf bifurcation), a small-amplitude limit cycle emerges. *p*21 exhibits weak, delayed pulses, while PUMA remains negligible. The system shows transient oscillations consistent with a reversible arrest without full commitment.

For *S* = 12 (robust oscillations), the trajectories form larger and slower cycles. *p*21 pulses are strong and regular, whereas PUMA remains below its threshold. This configuration reflects a sustained arrest state, which may progressively evolve toward senescence if *p*21 exceeds higher cutoffs over time.

At *S* = 24 (extreme stress), the dynamics collapse onto a high fixed point as the feedback weakens (due to the reduction of *k*_cat_ at large *S*). In this regime, *p*21 expression is suppressed, while PUMA rises toward a high plateau, consistent with an early apoptotic commitment.

### 2.4. Oscillatory window and bifurcation landscape in the (S, Σ) plane

Figure 4 summarizes the dynamical structure of the *p*53–*Mdm*2–*p*21–PUMA network across the (*S*, Σ) plane. The maps combine steady and oscillatory readouts: the first row reports maximal concentrations of *p*53, *p*21, and PUMA, while the second row shows their one–sided amplitudes 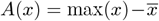 evaluated on a late-time window (60–72 h). Superimposed dotted and dash–dotted curves indicate the numerically determined Hopf loci 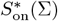 and 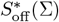, which delimit the region of sustained oscillations.

**Figure 4.**
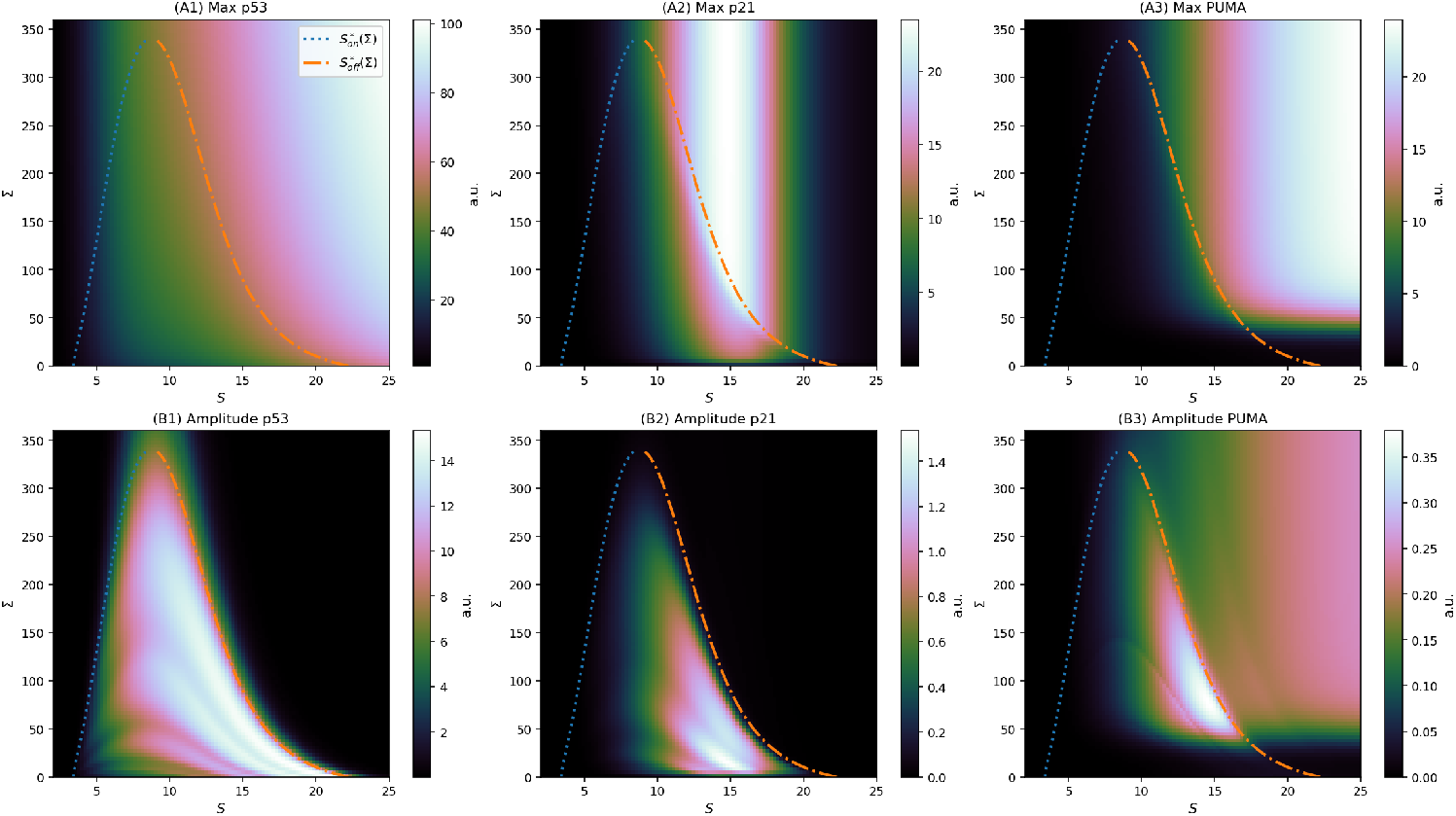
Oscillatory window and bifurcation landscape in the (*S*, Σ) plane. (A1–A3) Maximal concentrations and (B1–B3) one–sided amplitudes of *p*53, *p*21, and PUMA. Dotted and dash–dotted curves show the Hopf loci 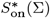 and 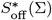. Increasing Σ shifts the oscillatory domain rightward and compresses it, eventually collapsing the pulses into a steady regime.

As Σ increases, the onset 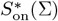 (dotted) shifts rightward and the offset 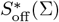 (dash–dotted) moves leftward, progressively narrowing the oscillatory band. For Σ ≳ 3.2 × 10^2^, the two boundaries nearly coincide and the system becomes aperiodic for all *S*. This behaviour reflects a decreasing effective loop gain in the *p*53→*Mdm*2 feedback: STAT1 enters through the regulatory factor 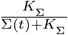 in the *Mdm*2_RNA_ equation, so increasing Σ reduces *Mdm*2 production and damps the feedback. At low Σ, moderate stress destabilises the fixed point (Hopf onset), whereas at higher *S* the stress–dependent term *k*_cat_(*S*) decreases the *Mdm*2-mediated p53 degradation rate, restoring stability (Hopf offset). Together, these mechanisms explain the closed oscillatory window bounded by 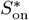 and 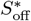. Figure S2 in the supplement illustrates Σ effect on bifurcation values.

Maximal concentrations (A1–A3). *p*53 reaches its highest levels along the upper-right flank of the Hopf region, reflecting the cumulative accumulation under sustained stress. *p*21 displays a more selective pattern: high expression is confined to a narrow intermediate-*S* domain due to the combined stress gating 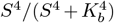 and the biphasic modulator *f*_hill_(*S*). PUMA, in contrast, increases almost monotonically with both *S* and Σ, dominated by the Σ-dependent co-activation *κ*_1_(Σ); its behaviour is only weakly influenced by the Hopf contours.

Oscillation amplitudes (B1–B3). *p*53 amplitude rises sharply at 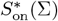, peaks inside the Hopf domain, and vanishes at 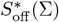. *p*21 follows a similar but narrower ridge and decays more rapidly with Σ, mirroring the attenuation of the upstream feedback. PUMA exhibits only minor oscillatory modulation, confirming that it integrates the steady (non-pulsatile) component of the *p*53 signal.

The combined analysis delineates stress–response zones in the (*S*, Σ) plane. Moderate *S* and intermediate–high Σ values place cells near the *p*53*/p*21 amplitude ridge—favouring adaptive or senescent responses—whereas large *S* or Σ drive PUMA accumulation and suppress oscillations, biasing toward apoptosis. Thus, STAT1 acts as a bifurcation modulator that reshapes the oscillatory landscape of the *p*53 network, bridging dynamical stability with cell-fate transitions.

In addition to the stress level *S*, the model also exhibits bifurcation behaviour with respect to two other parameters: the saturation constant *K*_0_ controlling Mdm2-mediated degradation of p53, and the degradation rate *δ*_*M*_ of Mdm2. As shown in Fig. S4 (Supplementary Information), both parameters can delimit regions where Hopf bifurcations occur. Importantly, increasing STAT1 levels Σ modulates the extent of these bifurcation zones, but the effect is less pronounced than for the primary bifurcation parameter *S*. This indicates that while *K*_0_ and *δ*_*M*_ shape the stability boundaries of the p53 oscillator, the dominant control of oscillatory dynamics remains exerted by the stress input. A 3D version for Figure 4 is given in the supplementary Fig. S3.

Together, these results establish the dynamical consistency of the model. The *p*53–Mdm2 core admits a *single* equilibrium for any fixed stress level *S*, ruling out bistability and ensuring a history–independent response. As *S* increases, this equilibrium successively loses and regains stability through a pair of Hopf bifurcations that delimit a well-defined oscillatory window. Within this window, the feedback loop generates self-sustained *p*53 pulses whose amplitude and frequency are modulated by both stress and STAT1 levels. Beyond it, trajectories converge either to a low basal state (weak stress) or to a high steady state (extreme stress), corresponding respectively to physiological homeostasis and apoptotic commitment. Hence, the analysis of stability and bifurcation provides a rigorous foundation for interpreting the transition from stable to oscillatory regimes and, ultimately, for linking dynamical behaviour to cell-fate outcomes.

## 3. Cell-fate decision: predicting apoptosis or cell-cycle arrest from the model

In complex signalling systems such as the p53–Mdm2–p21–PUMA network, cellular outcomes emerge from the interplay of multiple regulatory inputs. Here, both the genotoxic stress level (*S*) and the STAT1 activity (Σ) act as upstream modulators, dynamically shaping the p53 response. While qualitative patterns (e.g., oscillatory versus steady behaviour) can often be related to stress intensity, predicting the ultimate cell fate—whether adaptive, senescent, or apoptotic—remains challenging when two or more control axes interact. As illustrated below, even for fixed parameter sets, distinct combinations of (*S*, Σ(*t*)) may yield temporally complex *p*21 and *PUMA* profiles whose shape alone does not immediately reveal the underlying decision. This motivates the development of a formal, quantitative *decision criterion* based on measurable dynamical thresholds, enabling systematic classification of single-cell trajectories into distinct fate domains.

We define a *generic cell-fate criterion under stress*, applicable to any single cell whether stress *S* and STAT1 activity Σ are constant or time-varying. In other words, the rule applies equally to arbitrary trajectories *S*(*t*) and Σ(*t*) and to the particular cases *S*(*t*) ≡ *S* and/or Σ(*t*) ≡ Σ. Several studies have attempted to extract fate predictors from p53 dynamics, relating pulsatile versus sustained responses to transient arrest, senescence, or apoptosis (Lahav et al., 2004; Geva-Zatorsky et al., 2006; Batchelor et al., 2011; Purvis et al., 2012; Stewart-Ornstein & Lahav, 2017). Quantitative descriptors such as the number of p53 pulses *n*_p53_, their amplitude *A*_p53_, or the cumulative exposure 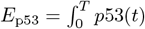 have been used to classify outcomes (Purvis et al., 2012). Downstream markers provided complementary criteria: *p*21 thresholds *θ*_21_ or time-integrated levels 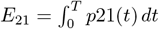 were associated with arrest and senescence (El-Deiry et al., 1993; Xiong et al., 1993; Abbas & Dutta, 2009; Barr et al., 2017; Hanson et al., 2019), while strong and sustained PUMA induction signalled apoptotic commitment (Jeffers et al., 2003; Nakano & Vousden, 2001; Sun et al., 2009). More integrative formulations compared these effectors directly, using ratios such as *R*_21_ = ⟨*p*21⟩_late_*/* max_*t*_ *p*21(*t*) (Stewart-Ornstein & Lahav, 2017) or *ρ* = ⟨*PUMA*⟩ */* ⟨*p*21⟩ (Kracikova et al., 2013), where large *ρ* values indicate apoptosis and small ones stable arrest. Although these indices capture important qualitative trends, they remain one-dimensional and context-specific. In contrast, our criterion introduces a two-dimensional and temporally resolved mapping from (⟨*p*21⟩, ⟨*PUMA*⟩) to discrete fate domains, defined by quantitative thresholds *µ*_21_ and *µ*_PU_ evaluated over early and stationary windows. This formulation unifies dynamic, molecular, and bifurcation-based perspectives into a continuous decision landscape that can be applied to heterogeneous cell populations and compared across archetypes.

1. *Early decision (before 60 h)*. We identify stable activation patterns that persist for at least the minimal required duration. If PUMA remains *continuously high* while p21 stays *below* its upper senescence threshold over a given time 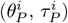, apoptosis will be triggered. If *p*21 lies in the *arrest band* (between 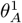 and 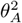) with PUMA 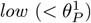 for 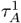, the cell cycle will be halted. If *p*21 lies in the *senescence band* (between 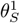 and 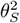) with PUMA *low* for 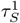, or is *stably very high* 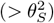 for 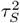, the cell will enter senescence.
2. *Early index (PUMA–p21 overlap)*. When both PUMA and p21 rise together, we look for a time window 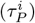 where *PUMA stays above its early threshold* and *p21 exceeds at least one of its activation thresholds*. Within that window we compute a balance index comparing peak PUMA to the sum of both peaks:

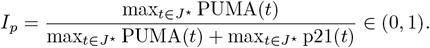 We then classify using two global cutoffs 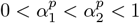:

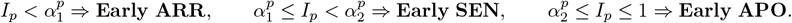

*Cycling is excluded here, since both pathways are already engaged*. If no qualifying window exists, no early decision is taken.
3. *Stationary decision (60–72 h)*. At the end of the experiment we average PUMA(⟨*PUMA*⟩) and p21(⟨*p*21⟩) over the late window (60–72 h) and compare them to global thresholds. If *both are low* (below *µ*_21_ and *µ*_PU_), cell cycle will continue. If *p21 is very high* (above *µ*_*S*_) with *PUMA low*, cell enters in senescence. If *p21 exceeds its decision level* (≥*µ*_21_) with *PUMA low*, cell cycle will be halted. If *PUMA is high* (≥*µ*_PU_) while *p21 remains below its decision level*, apoptosis will be triggered.
4. *Stationary index (both high late)*. If, in the late window, *both* PUMA and p21 are *high* (≥*µ*_PU_, *µ*_21_), we reuse a balance index based on late averages:

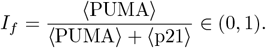

We classify using the same two global cutoffs:

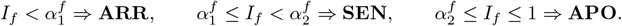

*Cycling is excluded since both effectors are above activation thresholds*.
5. *Effective decision (severity-first)*. Finally, we combine the early and stationary decisions. If both are defined, we report the most severe outcome: Return the effective fate

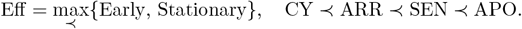

If only one decision is available, we report that one. Table 6 summarizes our fate criterion, a more formal version of the criterion is given in the supplement Box 1. To simulate fate decision for a generic cell, the thresholds are set from experimental dynamics and modelling practice. Typically between 50–80% of simulated maxima Geva-Zatorsky et al. (2006); Batchelor et al. (2008); Becker et al. (2010); Barr et al. (2017). Early thresholds are placed around 50–60%, final senescence higher (70–80%), then fine-tuned on simulated profiles for robustness and biological plausibility Ciliberto et al. (2005); Purvis et al. (2012). Throughout the criterion, *J* (*τ* ) denotes a sliding time window of duration *τ* over which activation or persistence is evaluated; for instance, 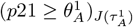 means That *p*21 remains above its threshold 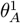 for at least 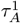 consecutive hours. When both PUMA and p21 are simultaneously active, we define *J* ^*\**^ as the maximal overlapping interval where their joint activation conditions hold, that is,

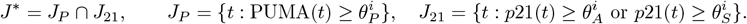

The balance indices *I*_*p*_ and *I*_*f*_ are then computed over *J* ^*\**^ and over the late window [60, 72] h, respectively. Global cutoffs *α*_*f*_ (and analogously *α*_*p*_) are fixed once by bounding late-window maxima with 95th percentiles across stresses, deriving an admissible interval [*I*^min^, *I*^max^], and choosing cutoffs by a small-margin heuristic or a minimax search.

### 3.1. Simulations for single cells

In this section, we apply the threshold-based decision criterion illustrated in Figure 6 to (*generic*) single-cell simulations that reproduce the behaviours summarized in Table 1. Besides constant stress and STAT1 levels, we also consider time-dependent inputs for *S*(*t*) and Σ(*t*). The profiles used (with their biological meaning) are listed in Table 3.

**Table 3.** Representative input profiles used to model stress *S*(*t*), active STAT1 (Σ(*t*)), or both, with corresponding mathematical expressions and biological/experimental interpretation.

| Profile | Applied to | Mathematical expression | Biological or experimental meaning |
| --- | --- | --- | --- |
| **Decreasing (exponential)** | Stress only | $S(t) = S_{\max} e^{-kt}$,<br>$k = \frac{\ln(S_{\max}/S_{\min})}{T_{\max}}$ | Models transient stresses with rapid decay, such as acute irradiation or oxidative stress, typically resolved by cellular repair mechanisms. |
| **Peak (Gaussian)** | Stress only | $S(t) = S_{\max} \exp\left(-\frac{(t - t_p)^2}{2\sigma^2}\right)$ | Short, high-intensity stresses such as bolus drug administration or heat shock pulses. |
| **Shifted (asymmetric) peak** | STAT1 ($\Sigma$) only | $\Sigma(t) = A(1 - e^{-\lambda_{\text{rise}} t}) e^{-\lambda_{\text{decay}} t}$ | Typical JAK–STAT kinetics: delayed activation and gradual decay driven by negative feedback and dephosphorylation. |
| **Sinusoidal (periodic)** | Both (experimental models) | $S(t) = \frac{S_{\max}}{2} \left[1 + \sin\left(\frac{2\pi t}{T_S}\right)\right]$<br>$\Sigma(t)$ analogously | Periodic input profile: circadian rhythms, fractionated treatment cycles, or simulated pulsed environmental and hormonal cues. |
| **Pulsed (repeated)** | Both (experimental/clinical) | $S(t) = \begin{cases} S_{\max}, & t \in [nT, nT + \tau_p) \\ 0, & \text{otherwise} \end{cases}$,<br>$\Sigma(t)$ analogously | Brief and regular pulses: repeated IFN injections, fractionated radiotherapy or chemotherapy, and cyclic immune signaling. |
| **Increasing (exponential)** | Both (mainly chronic stress/inflammation) | $S(t) = S_{\max}(1 - e^{-kt})$<br>$\Sigma(t) = \Sigma_{\max}(1 - e^{-kt})$ | Chronic or gradually rising inputs: progressive exposure to hypoxia, sustained DNA damage, or prolonged IFN signaling in inflammatory contexts. |

**Figure 5.**
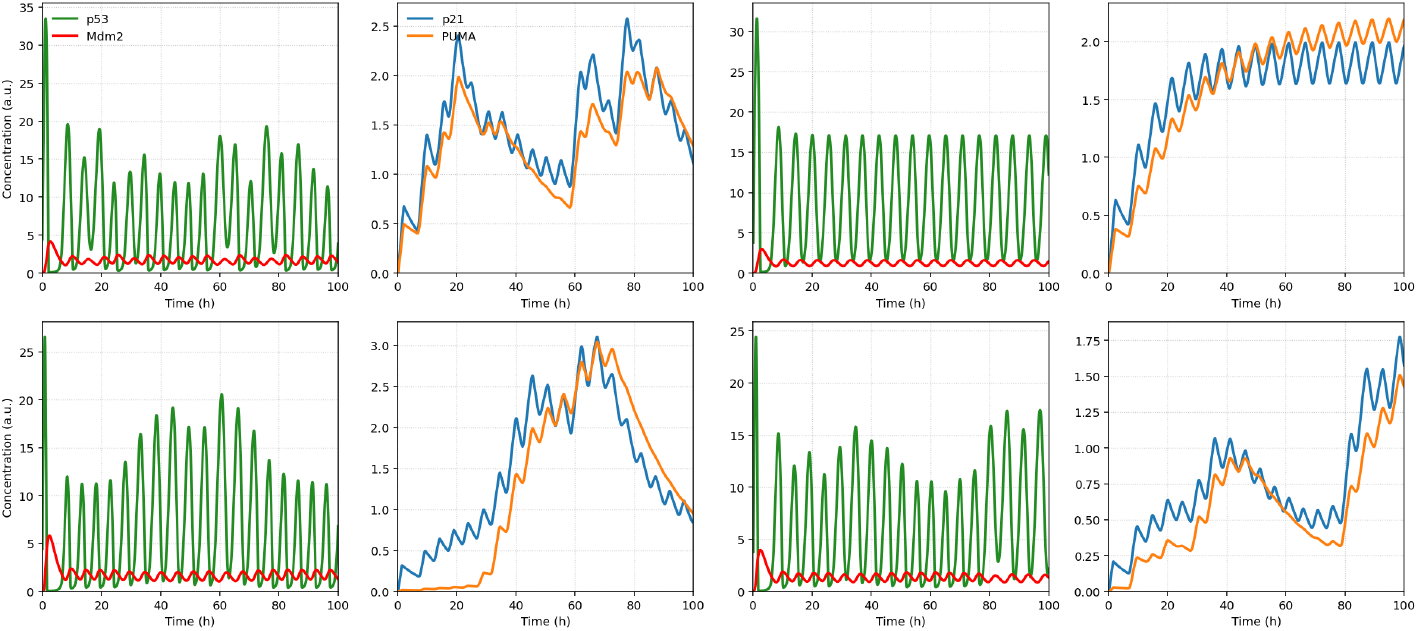
Paired single-cell simulations of the p53–Mdm2–p21–PUMA network under heterogeneous inputs. Each row corresponds to a distinct input scenario combining a fixed stress level (*S*) with either constant or time-varying STAT1 activity (Σ(*t*)). Odd columns display the dynamics of the core regulatory loop (*p*53, green; Mdm2, red), while the adjacent even columns show the corresponding downstream effectors (*p*21, blue; *PUMA*, orange). The diversity of oscillatory and transient behaviours illustrates how similar upstream conditions can produce distinct dynamical signatures, underscoring the need for a quantitative, threshold-based criterion to decode cell-fate outcomes.

**Figure 6.**
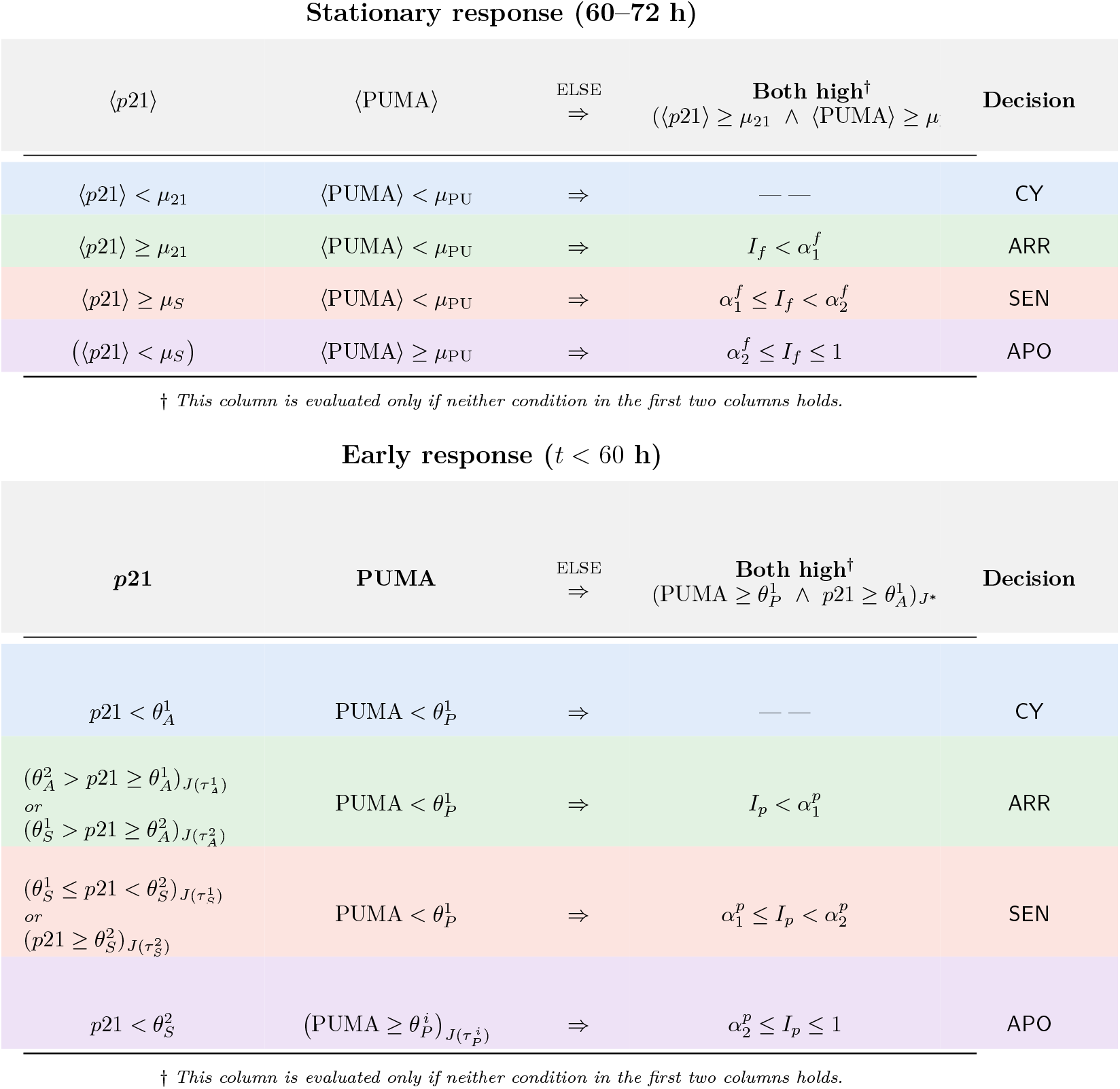
Decision rule with explicit gating: the *Both high* column is evaluated only if neither of the first two column conditions applies. Top: stationary means on [60, 72] h with *I*_*f*_ = ⟨PUMA⟩ */*(⟨*p*21⟩ + ⟨PUMA⟩). Bottom: early rule on sliding windows *J*(*τ*) with *I*_*p*_ = PUMA*/*(*p*21 + PUMA).

*Example combinations for simulations. S*(*t*) and Σ(*t*) are treated as independent inputs combined to emulate common experimental settings:

1. *Decreasing stress + offset STAT1 peak:* acute irradiation (fast DNA repair ⇒ *S* decays) followed by a delayed IFN-I/STAT1 activation through cGAS–STING signaling Brzostek-Racine et al. (2011); Yu et al. (2015); Feng et al. (2020).
2. *Pulsed stress + pulsed STAT1:* repeated or fractionated *S*(*t*) and bolus-like Σ(*t*) (e.g. IFN dosing). *Use case:* fractionated radiotherapy or chemotherapy combined with IFN-*α/γ* treatment Heitmeier et al. (1999); Giovannozzi et al. (2020); Kalliara et al. (2022).
3. *Slowly varying stress + tonic STAT1:* slowly increasing or periodic *S*(*t*) (chronic stress) combined with a tonic Σ(*t*) baseline (chronic inflammation) subject to slow attenuation. *Use case:* inflammatory microenvironments providing a persistent STAT1 background on which stress fluctuates Marié et al. (2021); Giles et al. (2017).

#### Box 1.

**Global** — Decision thresholds (stress-invariant)

**Early PUMA (APO):**

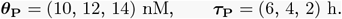

**Early** *p*21 **bands:**

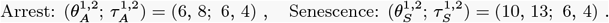

**Stationary gates (mean on** [60, 72] **h):**

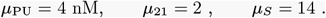

**Index cutoffs (for** *I*_*p*_ **and** *I*_*f*_ ):

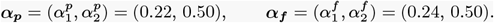

#### Cell fate under time-varying STAT1

To isolate the effect of STAT1, we fixed the stress level at a single value (*S* = 17) and varied *only* the STAT1 signal Σ(*t*).

We considered four representative temporal profiles—exponential decay, rising ramp, offset (asymmetric) peak, and sinusoid—each evaluated at two maximum amplitudes (Σ_max_ ∈ {50, 120}). All other parameters and initial conditions were kept identical. For each condition we simulated the *p*53–*Mdm*2 module together with the effectors *p*21 and PUMA, then applied our fate criterion. When Σ_max_ = 50 (rows A–D): In the *p*53–Mdm2 module, oscillations are sustained but of moderate amplitude for the decreasing and sinusoidal inputs, while the increasing profile drives a slow damping toward steady-state levels—consistent with irreversible senescence. The system remains in a low-STAT1 regime where PUMA activation is limited. No early apoptotic commitment occurs; instead, *p*21 dominates the response. Depending on the input waveform, the cell either stabilizes in a moderate-amplitude arrested state or gradually transitions toward a senescent plateau.

When Σ_max_ = 120 (rows E–H): Across these conditions, *p*53–Mdm2 oscillations are more strongly damped and settle around higher mean values, reflecting enhanced stabilization of nuclear p53. Elevated STAT1 strengthens PUMA induction, pushing trajectories toward apoptosis. In all cases, PUMA surpasses its apoptotic threshold early, although the timing and sharpness of commitment depend on waveform shape. The decreasing and asymmetric-peak profiles trigger the fastest apoptotic responses (*t* ∼ 20–25 h), whereas the sinusoidal input produces alternating windows of susceptibility: in-phase peaks with stress favor apoptosis, while out-of-phase segments allow transient arrest or delayed senescence.

Overall, these comparisons highlight that the *temporal architecture* of the STAT1 signal—its rise and decay kinetics, duration, and frequency—can substantially reshape fate outcomes even under the same stress intensity. By synchronizing (or desynchronizing) STAT1 peaks with *p*53 pulses, cells may shift between arrest, senescence, and apoptosis despite identical stress inputs.

#### Cell fate under time-varying STAT1 and stress

We vary *both* inputs in time and present the simulations in two complementary grids, keeping the paired layout (left: *p*53–Mdm2 with scaled *S*(*t*) and Σ(*t*); right: *p*21–PUMA with the early band and the stationary window [60, 72] h). Below is a detailed reading of Figure 8

**Figure 7.**
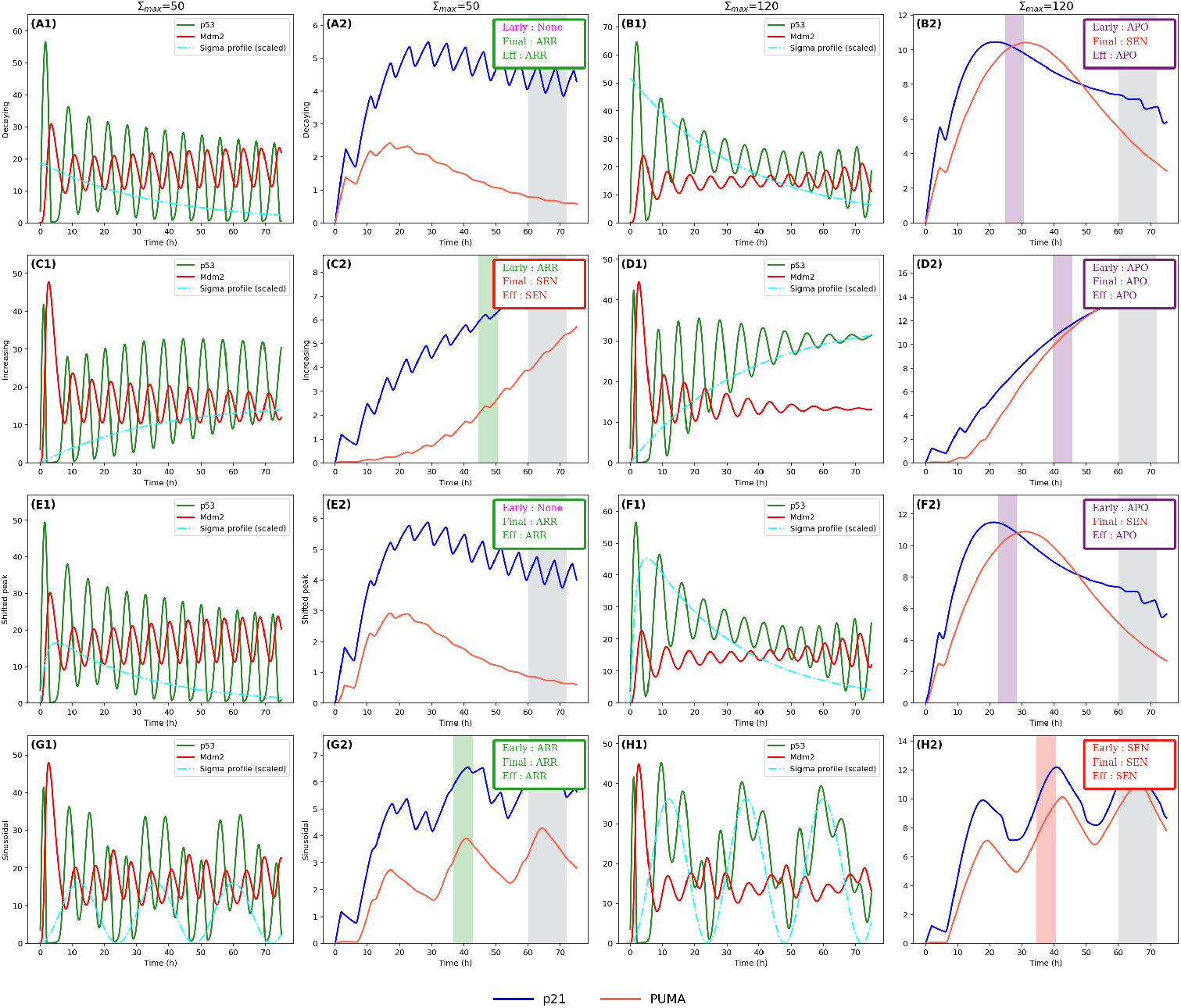
Impact of STAT1 waveform shape and amplitude on the p53–Mdm2–p21–PUMA dynamics at fixed stress (*S* = 17). Each row corresponds to a distinct STAT1 input profile (decreasing, increasing, shifted peak, sinusoidal), while paired columns show the *p*53–Mdm2 module (left) and downstream effectors *p*21 and *PUMA* (right). The upper half (Σ_max_ = 50) represents weak STAT1 activation, leading to arrest or senescence dominated by *p*21. The lower half (Σ_max_ = 120) illustrates strong STAT1 signaling, where PUMA surpasses its apoptotic threshold and early commitment to apoptosis occurs. Colored shaded bands mark the decision windows (early vs. stationary). Together, these simulations demonstrate that not only the amplitude but also the *temporal shape* of the STAT1 signal—its rise, decay, and oscillatory phase—critically modulates fate determination under the same stress intensity.

**Figure 8.**
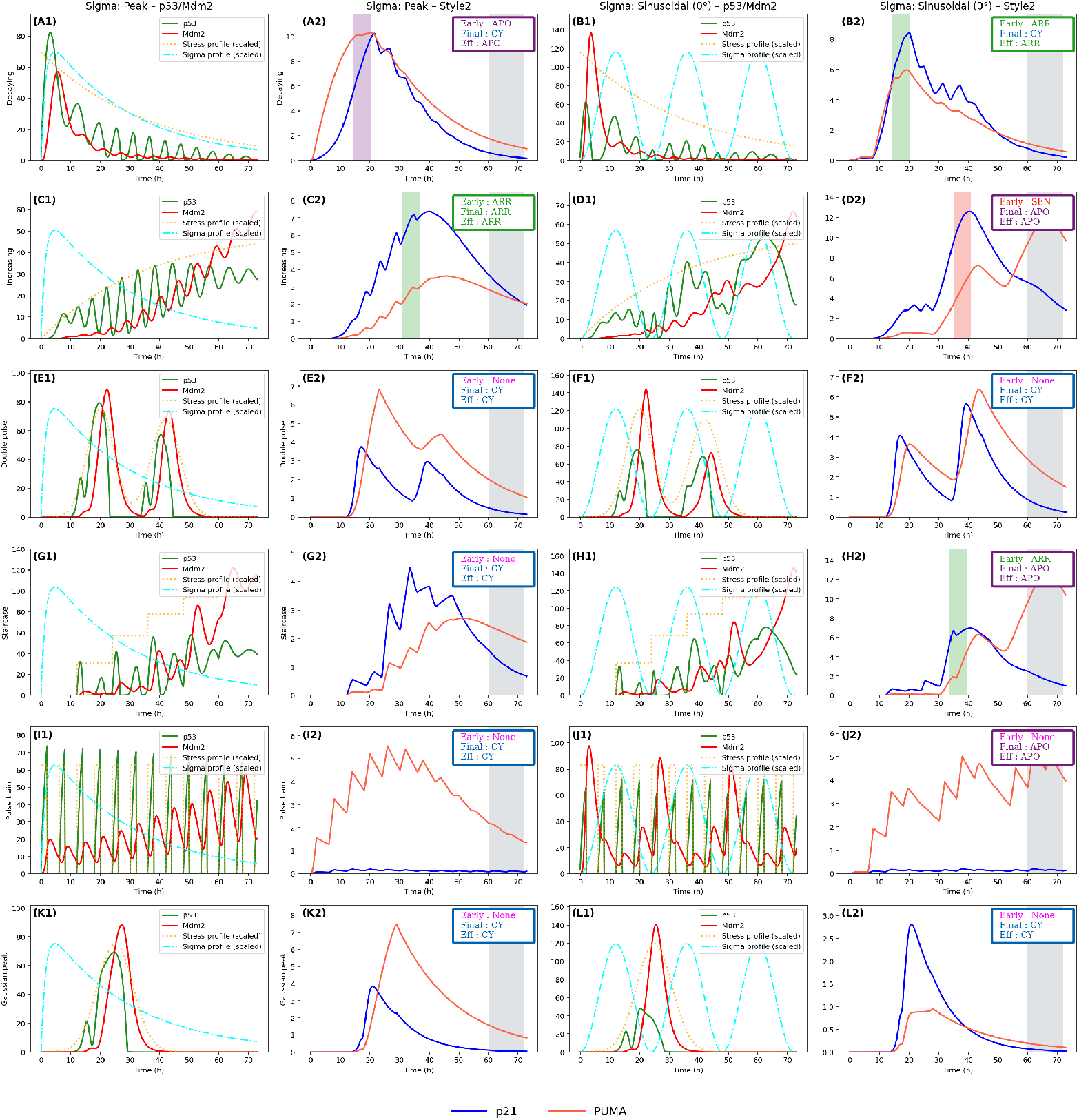
Coupled profiles *S*(*t*) and Σ(*t*).

*Row A — Decaying S*(*t*). (A1) Σ(*t*) has an offset peak while *S*(*t*) decays, yielding initially large but damped *p*53 pulses. (A2) Early APO appears when the first overlap is strong, but PUMA later falls and the [60, 72] h average stays sub-plateau, giving CY at stationarity (APO → CY by lack of duration).

*Row B — Decaying S*(*t*) *with in-phase sinusoidal* Σ(*t*). (B1) Sinusoid crests arrive as *S*(*t*) is already declining, fragmenting the duty-cycle of *p*53. (B2) Early ARR from *p*21 and ARR (or CY) at stationarity; phase cannot compensate for the loss of duration caused by decaying *S*(*t*).

*Row C — Increasing S*(*t*).. (C1) Pulses of *p*53 lengthen gradually as *S*(*t*) rises. (C2) The single Σ peak occurs too early, mainly feeding *p*21: early ARR, and ARR (or CY) at stationarity because PUMA is insufficient within [60, 72] h.

*Row D — Increasing S*(*t*) *with in-phase sinusoidal* Σ(*t*). (D1) A well-timed Σ crest on high *S*(*t*) produces a long *p*53 pulse near the stationary window. (D2) Early SEN (or ARR depending on thresholds) followed by APO final; constructive overlap in [60, 72] h carries PUMA across the plateau (SEN → APO).

*Row E — Double-pulse S*(*t*).. (E1) Two stress bursts; the Σ peak aligns mostly with the first. (E2) No early band and CY final: the second pulse lacks STAT1 assistance, so the useful PUMA area remains too small.

*Row F — Double-pulse S*(*t*) *with in-phase sinusoidal* Σ(*t*). (F1) Several partial overlaps; some *p*53 peaks are tall but short-lived. (F2) CY final despite decent PUMA maxima: the [60, 72] h mean stays below *µ*_PU_. Duration beats instantaneous amplitude.

*Row G — Staircase S*(*t*). (G1) Each plateau lengthens *p*53 pulses; the Σ peak is early and then decays. (G2) No early band and CY final: staircase growth without a late Σ assist is insufficient for PUMA in the stationary window.

*Row H — Staircase S*(*t*) *with in-phase sinusoidal* Σ(*t*). (H1) A Σ crest on a high plateau yields a very long *p*53 pulse just before stationarity. (H2) Early ARR from *p*21, then APO final: PUMA crosses the plateau within [60, 72] h (ARR → APO).

*Row I — Pulse train S*(*t*). (I1) Many narrow *p*53 pulses; the single Σ peak covers only a fraction. (I2) No early band and CY final: the useful integral is too small despite repeated spikes.

*Row J — Pulse train S*(*t*) *with in-phase sinusoidal* Σ(*t*). (J1) Repeated short overlaps filtered by the *p*53–Mdm2 loop. (J2) No early band but APO final: a sufficiently long late overlap occurs inside [60, 72] h and lifts PUMA above threshold.

*Row K — Single Gaussian S*(*t*). (K1) Very strong but brief stress; Σ then decays. (K2) No early band and CY final: large PUMA peak but insufficient duration for the stationary mean.

*Row L — Single Gaussian S*(*t*) *with in-phase sinusoidal* Σ(*t*). (L1) Alignment can extend one *p*53 pulse, yet not long enough within [60, 72] h. (L2) No early band and CY final: confirms that, for an isolated pulse, the mean over the stationary window is decisive.

The common motifs throughout the grid are: (i) Duration inside the stationary window dominates, not the instantaneous maxima. (ii) Phase matters: constructive overlap near [60, 72] h drives APO even when no early band is present. (iii) Asymmetry between PUMA and *p*21: *p*21 accumulates and often yields early ARR/SEN, whereas PUMA requires long, well-timed *p*53 pulses, hence frequent early–final mismatches. (iv) The *p*53–Mdm2 loop behaves as a low-pass filter; fast or sparse overlaps are attenuated.

All full time courses for time varying stress or Stat1 profiles and different initial conditions for *p*21 and PUMA are available in Supplementary Section S4 on the Figure S5,Figure S6 and Figure S7.

## 4. From single-cell rules to population outcomes (via archetypes)

### 4.1. From single cells to archetypes

Our goal is to test whether the threshold–based fate criterion, validated on a generic single cell, also explains outcomes at the *population* level—even when different populations display lineage–specific behaviours. To do so, we apply the single–cell decision rule of Fig. 6 at population scale: each cell is classified independently with the same severity–first priority CY ≺ ARR ≺ SEN ≺ APO. Population fractions are simple averages across cells, and uncertainty is reported by nonparametric bootstrap.

We represent a cell population by an *archetype* 𝒜 = (Θ, 𝒟_*S*_, 𝒟_Σ_, ℋ). Here, Θ denotes the ODE parameter set; 𝒟_*S*_ and 𝒟_Σ_ specify the stress and STAT1 inputs (constants or sampled distributions, e.g. log–normal micro–stress or STAT1 dispersion); and ℋ collects the decision thresholds used by the rule in Fig. 6 (early and stationary cut–offs). Differences between populations are therefore encoded by changes in Θ, in the input statistics (𝒟_*S*_, 𝒟_Σ_), and—when distinct sensitivities are required—by archetype–specific thresholds ℋ; the dynamical equations are not modified.

Single–cell heterogeneity within an archetype is introduced by drawing, for each cell, a parameter vector from independent log–normal laws centred on the archetype “Center” values in Table 5 with the listed coefficients of variation (CV). For any parameter *p* with mean *µ*_*p*_ and CV *c*_*p*_,

**Table 4.** Representative population outcomes ∼48–72 h after single-dose IR. Percentages are approximate unless stated.

| Cell line (p53) | Gy | $S$ | Cycle (%) | Arrest (%) | Senescence (%) | Apoptosis (%) |
| --- | --- | --- | --- | --- | --- | --- |
| HCT116 (WT)<sup>a</sup> | 0 | 0 | $\gtrsim 80$ | n.d. | $\approx 0$ | 5–10 |
| | 2 | 3 | n.d. | n.d. | n.d. | $\approx 24$ |
| | 5 | 6.5 | n.d. | ↑ | low | 10–20 |
|  | 10 | 10 | n.d. | n.d. | low–mod. | 40–70 |
| MCF-7 (WT, casp-3<sup>-</sup>)<sup>b</sup> | 0 | 0 | $\gtrsim 85$ | $\sim 10$ | $\approx 0$ | $< 5$ |
| | 4 | 3.5 | ↓ | mod. | ↑ | $< 10$ |
|  | 8–10 | 10 | low | mod. | **high** | low–mod. |
| | 20 | 25 | $\approx 0$ | – | **very high** | low |
| U2OS (WT)<sup>c</sup> | 0 | 0 | $\gtrsim 85$ | $\sim 10$ | $\approx 0$ | $< 5$ |
|  | 6 | 8 | n.d. | mod. | up to 50–60 | low |
| | 10 | 10 | n.d. | mod. | mod. | $\approx 32$ |

<sup>a</sup> HCT116: apoptosis measurable after IR (e.g., visualization and quantification frameworks) and early APO $\sim 10$ –20% around 5 Gy Bae et al. (2018); p53-driven fate engagement at higher doses Purvis et al. (2012).
<sup>b</sup> MCF-7: senescence-dominant from 8–10 Gy; very high SEN at 20 Gy Bravatà et al. (2015).
<sup>c</sup> U2OS: representative apoptosis fractions reported at 10 Gy (72 h) in radioresponse contexts ?.

**Table 5.** Randomly sampled parameters: central value and CV, by archetype.

| Parameter (sampled) | HCT116-like |  | MCF-7-like (casp-3<sup>-</sup>) |  | U2OS-like |  |
| --- | --- | --- | --- | --- | --- | --- |
|  | Center | CV | Center | CV | Center | CV |
| $\beta_{\max}$ | 60 | 0.20 | 60 | 0.20 | 60 | 0.18 |
| $\hat{\beta}_{p21}$ | 0.02 | 0.20 | 0.80 | 0.20 | 0.06 | 0.18 |
| $\varphi_{pu}$ | 1.70 | 0.25 | 0.12 | 0.25 | 5.60 | 0.18 |
| $\varphi_{p21}$ | 13.8 | 0.25 | 14.8 | 0.25 | 10.5 | 0.18 |
| $\delta_p$ | 0.50 | 0.15 | 0.50 | 0.15 | 0.50 | 0.12 |
| $\delta_{p21}$ | 0.10 | 0.15 | 0.10 | 0.15 | 0.10 | 0.12 |
| $\delta_{pu}$ | 0.05 | 0.15 | 0.05 | 0.15 | 0.05 | 0.12 |
| $\delta_m$ | 0.40 | 0.15 | 0.40 | 0.15 | 0.40 | 0.12 |
| $\delta_M$ | 0.40 | 0.15 | 0.40 | 0.15 | 0.40 | 0.12 |
| $K_a$ | 15 | 0.10 | 15 | 0.10 | 16.0 | 0.08 |
| $K_b$ | 8 | 0.10 | 8 | 0.10 | 8.0 | 0.08 |
| $K_{PUMA}$ | 9 | 0.10 | 9 | 0.10 | 9.0 | 0.08 |
| $K_{21}$ | 6.3 | 0.10 | 6.3 | 0.10 | 6.8 | 0.08 |
| $\Sigma$ (STAT1)<sup>a</sup> | 100 | 0.20 | 80 | 0.25 | 110 | 0.25 |
| $S$ (micro-mult.)<sup>b</sup> | 1.0 | 0.05 | 1.0 | 0.05 | 1.0 | 0.06 |
| IC $p21_0$<sup>c</sup> | 0.5 | 0.20 | 0.5 | 0.20 | 0.5 | 0.20 |
| IC $PUMA_0$<sup>c</sup> | 0.5 | 0.20 | 0.5 | 0.20 | 0.5 | 0.20 |

<sup>a</sup> Log-normal draw around the archetype mean (100/80/110).
<sup>b</sup> Log-normal multiplicative factor applied to baseline $S$ (micro-heterogeneity).
<sup>c</sup> LogN(mean = 0.5, CV = 0.20) draw, shifted by $-0.5$ and truncated at 0.
<sup>d</sup> Parameters correspond exactly to those used in the population simulations.

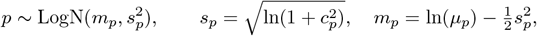

truncated to physiological bounds (±3*s*_*p*_). CV tiers reflect expected single–cell dispersion while maintaining oscillator robustness: 8–10% for affinity/threshold constants (*K*’s), 15% for decay/catalytic rates (12% in U2OS), and 20–25% for activation gains and basal terms. STAT1 activity Σ is drawn log–normally around an archetype–specific mean with CV 20–25%; a micro–stress multiplier on *S* uses a small CV (5–6%) to mimic dose dispersion; initial conditions for *p*21_0_ and PUMA_0_ are log–normal (mean 0.5, CV 20%), shifted and truncated at 0. Draws are independent; when comparing doses for the same cell, the sampled vector is held fixed to avoid Monte Carlo re–labelling. Archetype–specific decision thresholds ℋ are tuned once at *S* = 10 and then kept fixed for hold–out *S*.

#### Pre–registered constraints on the p53–Mdm2 core

To allow modest lineage–specific kinetics without over–parameterisation, each archetype permits only two bounded deviations in the p53–Mdm2 core: (i) the Mdm2 protein degradation rate *δ*_*M*_, and (ii) the transcription/translation delay ratio *k*_tr_*/k*_tl_. Both are restricted to ±10% around reference values, with the product *k*_tr_*k*_tl_ held constant to preserve the ∼ 6 h oscillation period. All other core constants (*K*_0_, *k*_*u*_, *K*_*u*_, *δ*_*p*_, …) are fixed.

#### Calibration and population readouts

We calibrated three archetypal lines (HCT116–like, MCF–7–like, U2OS–like) to match representative population outcomes ∼48–72 h after single–dose IR (Bae et al. (2018); Bravatà et al. (2015)). Calibration adjusts only lineage–specific central parameters and, when needed, the STAT1 co–activation strength and the threshold set ℋ; the zoned decision rule and all ODEs remain unchanged. For each archetype, fate fractions *f*_*X*_ (𝒜) are computed as averages of single–cell labels, and uncertainty is quantified by nonparametric bootstrap.

### 4.2. Population simulations

We simulate *N* =1500 cells per archetype and restrict *S* to literature-supported levels. We present here the results for the MCF-7–like archetype as introduced in Section S5. The same procedure is applied unchanged to the HCT116–like and U2OS–like archetypes; their complete single–cell panels, population scatters, fate fractions, and sensitivity checks are provided in Supplementary Figures Figure S9–Figure S11 and Supplementary Boxes 2-3. A compact comparison across all three archetypes is shown in Figure 11. The archetype analyses below reuse exactly the single-cell gates defined above; for each archetype, cutoffs and *I*_*f*_ (*I*_*p*_) partitions are held *stress-invariant* (no per-*S* threshold tuning).

**Figure 9.**
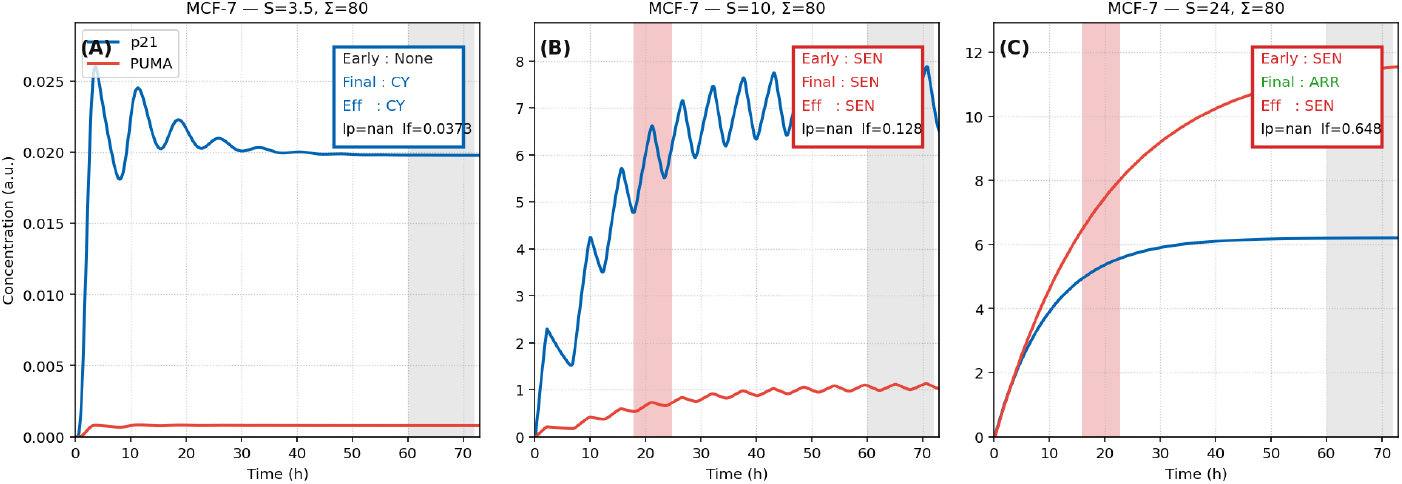
MCF-7 single-cell trajectories at Σ = 80 for three stress levels. Shaded red bands mark the first early-commitment window; the grey band is the stationary window (60–72 h). **(A)** *S* = 3.5: both effectors remain low; no early gate is crossed and the stationary call is cycling (CY). **(B)** *S* = 10: *p*21 pulses build and cross the senescence band while PUMA stays sub-threshold; the cell is classified SEN both early and late. This reflects the *p*21 branch dominating at moderate stress. **(C)** *S* = 24: PUMA rises to a high plateau and *p*21 settles at a mid/high level—its production attenuates at high stress in this MCF-7 parametrization—yielding an early SEN window and a SEN decision in the stationary phase.

**Figure 10.**
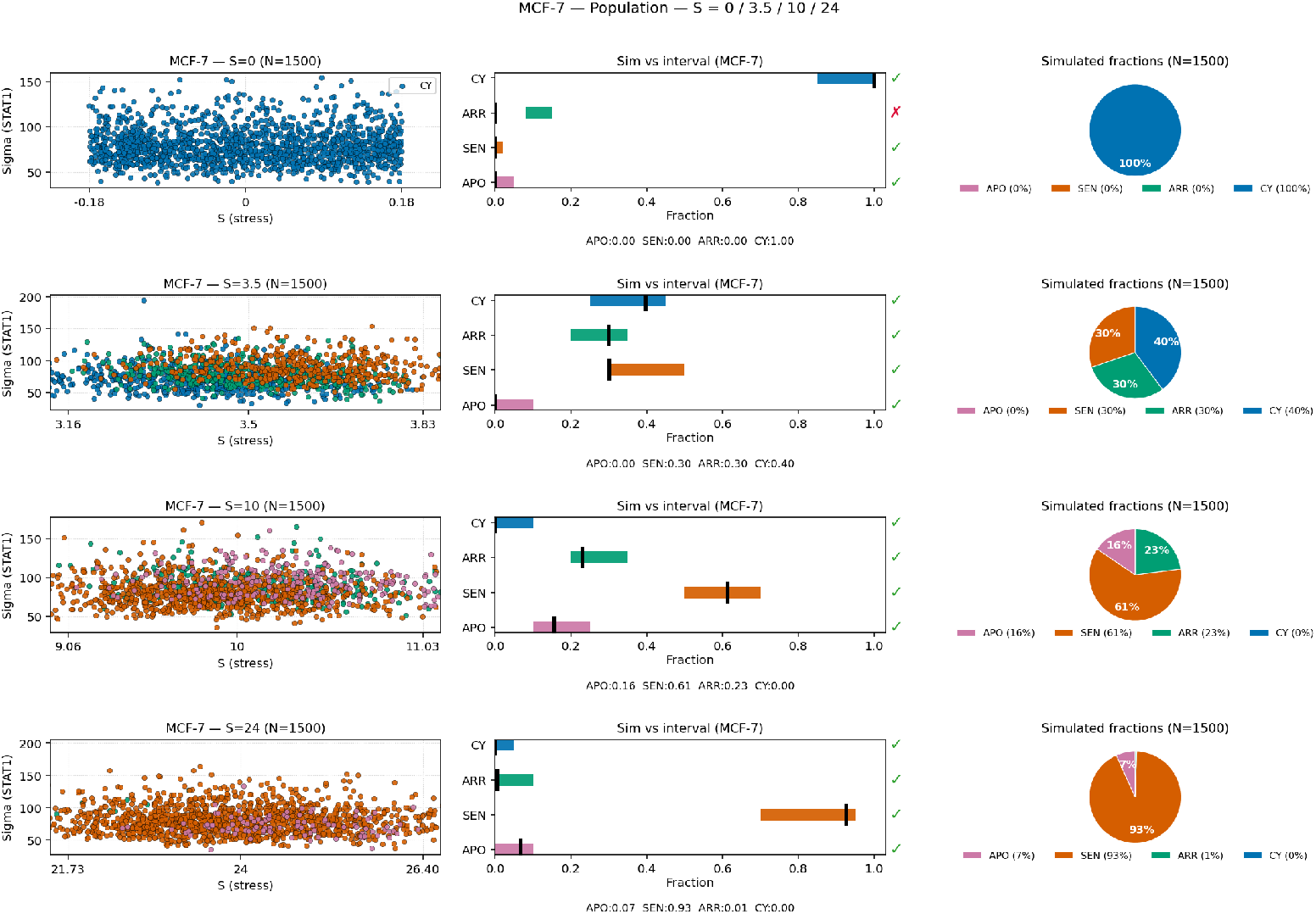
MCF-7-like simulation results for population). Population-level simulations for the MCF-7 archetype, showing the distribution of fates across cells.

**Figure 11.**
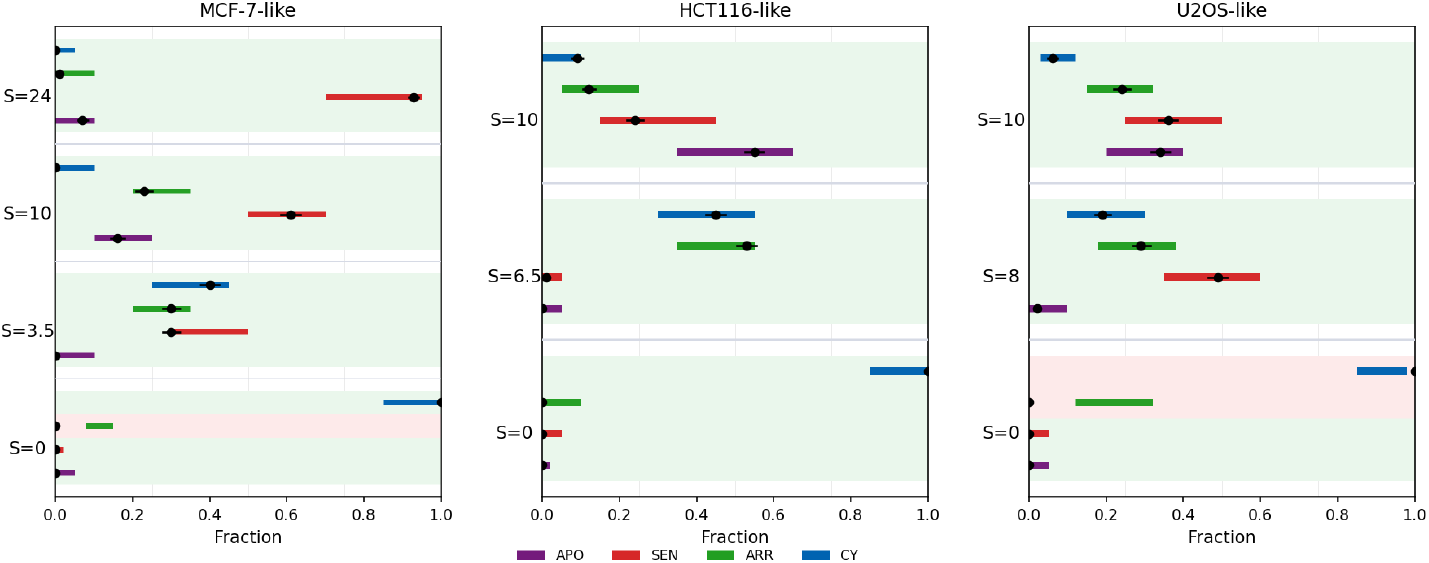
Summary of the 3 archetype simulations

### 4.3. MCF–7 archetype

We expect a small amount of arrest and apoptosis at very low stress, but above all massive senescence instead of apoptosis at extreme stress. The specific parameter values used for the MCF-7 archetype are listed in Table 6:

**Table 6.** MCF-7-specific parameters used for the population simulations (*N* =1500 cells per stress level). Σ denotes the baseline STAT1 activation level (log-normal dispersion CV = 0.25), while *S* is the DNA-damage stress input scanned in [0, 25]. All other constants are inherited from Table 5. Decision thresholds and *α* indices are given in the box bellow.

| Parameter | Value | Parameter | Value | Parameter | Value |
| --- | --- | --- | --- | --- | --- |
| *PUMA branch* |  |  |  |  |  |
| $\beta_{\text{PUMA}}$ | 0.60 | $\varphi_{\text{pu}}$ | 0.12 | $K_a$ | 15.0 |
| $K_{\text{PUMA}}$ | 9.0 | $\delta_{\text{pu}}$ | 0.05 | | |
| *p21 branch* |  |  |  |  |  |
| $\hat{\beta}_{21}$ | 0.80 | $\varphi_{21}$ | 14.8 | $K_b$ | 8.0 |
| $K_{21}$ | 6.3 | $\delta_{21}$ | 0.10 | | |
| *Stress and STAT1 context* |  |  |  |  |  |
| $\Sigma$ | 80 (baseline mean) | $S$ | varied on [0, 25] | | |

#### Box 2.

**MCF -7-like** — Decision thresholds (global)

**Early PUMA (APO):**

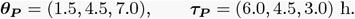

**Early** *p*21 **bands:**

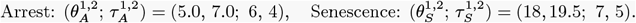

**Stationary gates (mean on** [60, 72] **h):**

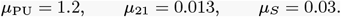

*µ*_PU_ = 1.2, *µ*_21_ = 0.013,

**Index cutoffs (used for** *I*_*p*_ **and** *I*_*f*_ ):

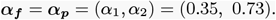

To reproduce the experimental MCF–7 sequence (cycling → arrest → senescence) without altering the ODE structure or the global thresholds, we used a *physiological zone calibration*. In this scheme, only context variables that biologically depend on the stress regime were adjusted: mean STAT1 activation (Σ), branch weights (*φ*_pu_, *φ*_21_), and the fraction of cells pre-activated in p21 (*seeding* fraction). Specifically, Σ was slightly increased at intermediate and high stress (*S*=10, 24) to mimic the known enhancement of JAK/STAT1 signaling under DNA damage; the PUMA branch was mildly attenuated while the p21 branch was reinforced at high stress, reflecting the experimentally observed bias toward senescence in MCF–7 cells. These modulations affect only the statistical initialization of the cell population and not the dynamical equations or the decision thresholds (*α*_1,2_, *θ, µ*), thereby preventing overfitting. The resulting configuration remains fully predictive within each stress zone and captures the physiological transition from mixed arrest/senescence at *S*=10 to irreversible senescence at *S*=24.

For each stress level *S* and fate *k* ∈ {CY, ARR, SEN, APO}, the literature specifies an admissible interval [*ℓ*_*S,k*_, *u*_*S,k*_]. From *N* =1500 simulated cells we estimate 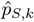, the empirical fraction per fate. On the right-hand panels, colored bars represent the literature intervals, and the black ticks show the simulated 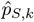; a green 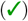 or red 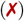 mark indicates whether 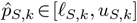.

#### MCF–7-like: why at high S we have SEN (threshold view)

We keep the ODEs unchanged and modulate the outcome solely through *global thresholds*. (i) Early APO remains rare because the PUMA gates are stringent: *θ*_PUMA_ = (1.5, 11.5, 12.0) with durations *τ*_PUMA_ = (6, 4.5, 3) h; short or modest PUMA transients fail to reach these levels. (ii) Late SEN is easy to attain: the stationary gates are *µ*_PUMA_ = 1.2, *µ*_21_ = 0.013, with a very low senescence trigger 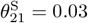; at high stress, p21 steadily surpasses this limit while PUMA remains below *µ*_PUMA_. (iii) When both markers coexist, the overlap index uses two global cutoffs *α* = (0.35, 0.73), so the typical p21 ≫ PUMA ratio is classified as SEN, with only clearly PUMA-dominant overlaps promoted to APO. Net effect: senescence becomes the dominant fate at large *S*, consistently reproducing the MCF–7 high-stress phenotype.

We assess, for each stress *S* 0, 3.5, 10, 24 and fate *k* ∈ {CY, ARR, SEN, APO}, the point estimate 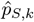 against the admissible literature interval [*ℓ*_*S,k*_, *u*_*S,k*_], using the coverage indicator *c*_*S,k*_, the normalized gap *g*_*S,k*_, and the binomial Wilson 95% confidence interval [lo_*S,k*_, hi_*S,k*_] as defined in Sec. S5.1.

*S*=3.5. Empirical fractions 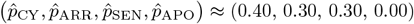 fall within their bands CY ∈ [0.25, 0.45], ARR ∈ [0.20, 0.35], SEN ∈ [0.30, 0.50], APO ∈ [0, 0.10]. Hence *c*_*S,k*_ = 1 and *g*_*S,k*_ = 0 for all *k*, giving *C*_3.5_ = 1.00 and *G*_3.5_ = 0. Wilson 95% CIs (*N* =1500) intersect the bands in all cases (e.g. for 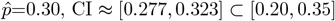; for 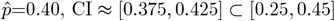; for 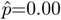, the Wilson upper bound ≲ 0.01 remains inside [0, 0.10]). Decision: **PASS**.

*S*=10. Empirical fractions ≈ (0.00, 0.23, 0.61, 0.16) for (CY, ARR, SEN, APO) lie inside the literature bands [0, 0.10], [0.20, 0.35], [0.50, 0.70], [0.10, 0.25], respectively. Thus *c*_*S,k*_=1 and *g*_*S,k*_=0 for all *k*, yielding *C*_10_=1.00 and *G*_10_=0. Wilson CIs overlap the bands in every class (e.g. 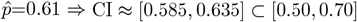). Decision: **PASS**.

*S*=24. Empirical fractions ≈ (0.00, 0.01, 0.93, 0.07) are within the target bands [0, 0.05], [0, 0.10], [0.70, 0.95], [0, 0.10]. Hence *c*_*S,k*_=1, *g*_*S,k*_=0, *C*_24_=1.00, *G*_24_=0. Wilson CIs remain inside/overlapping the bands (e.g. 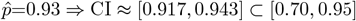). Decision: **PASS**.

*S*=0. At basal stress, the model predicts a fully cycling population (**CY** ≈ 1.00) with negligible activation of p53, p21, or PUMA. This outcome does not arise from missing literature bands but from the intrinsic structure of the model: at *S*=0, the transcriptional drives for p53 and its downstream targets vanish, preventing protein production and hence any irreversible response. In agreement with Fig. S7, when initial concentrations remain below the thresholds of irreversible fates (apoptosis or senescence), the system stays trapped in the cycling basin corresponding to its stress level. Accordingly, interval-based validation is not applicable here; we report 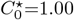 and 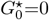 for the residual classes, acknowledging that the lack of activation reflects a physiological rather than statistical limitation of the model.

#### Summary

For MCF–7-like populations at *S*=3.5, 10, 24, we obtain full interval coverage with *C*_*S*_=1.00, *G*_*S*_=0, and Wilson 95% CIs overlapping the literature intervals in all classes, thus meeting the *PASS* criterion. At *S*=0, the available classes also pass 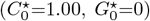, while CY is not band–validated due to missing literature range. These results confirm a consistent calibration across the MCF–7 stress sequence.

### 4.4. Population–level agreement for HCT116-like and U2OS–like

#### HCT116-like

Across the three stress points (*S*=0, 6.5, 10), the agreement between simulated and literature fractions is uniformly strong (*grit: 12/12*,✓). At *S*=0, the cohort remains fully cycling (*CY* ≈ 1.00) with negligible other classes; all bins lie within their admissible ranges, Wilson 95% confidence intervals intersect the bands, and global distances are minimal. At *S*=6.5, the population split (*ARR* ≈ *0.53, CY* ≈ *0.45, SEN* ≈ *0.01, APO* ≈ *0.00* ) fits squarely inside the literature intervals, yielding a non-significant goodness-of-fit and small classwise residuals. At *S*=10, the response shifts toward apoptosis and senescence (*APO* ≈ *0.55, SEN* ≈ *0.24, ARR* ≈ *0.12, CY* ≈ *0.09* ); all values fall within the expected ranges, with tight CIs and low normalized gaps. Overall, HCT116-like simulations display excellent concordance from mid to high stress without systematic deviations.

#### U2OS–like

Over the three evaluated levels (*S*=0, 8, 10), the overall agreement is high (*grid: 10/12* ✓). At *S*=0, the simulation yields a nearly pure cycling population (*CY* ≈ 1.00), whereas the literature includes a minor arrest fraction; this explains one of the two misses (*ARR*_*S*=0_ ) and reflects the absence of p53 activity at basal stress rather than a numerical error. At *S*=8, the simulated mix (*SEN* ≈ *0.49, ARR* ≈ *0.29, CY* ≈ *0.19, APO* ≈ *0.02* ) lies entirely within literature intervals with overlapping Wilson CIs and very small residuals, indicating near-perfect alignment. At *S*=10, the simulation slightly overestimates cycling (*CY* ≈ 0.06 vs. band upper bound ≈ 0.05), while *APO, SEN*, and *ARR* remain inside their respective ranges. This deviation is marginal and does not alter the fate hierarchy (*SEN* > *APO* > *ARR* > *CY* ). Global distances stay below 0.1 and all Wilson CIs intersect their bands.

Across archetypes, mid-stress points again yield the tightest SIM–LIT alignment. The two isolated deviations (*ARR*_*S*=0_ in U2OS, *CY*_*S*=0_ marginally above its band) are mechanistically interpretable: they reflect basal inactivity or residual proliferative potential rather than parameter mismatch. Elsewhere, every fate lies within its literature band with overlapping 95% Wilson confidence bounds and normalized gaps *G*_*S*_ ≤ 0.1, confirming excellent global concordance under stress-invariant gates.

## Discussion

Our stress–aware model couples the *p53–Mdm2* core to *p21* /*PUMA* and treats *STAT1* (Σ) as a modulatory input. In doing so, it recovers hallmark features of the DNA-damage response while adding a second, cytokine-controlled axis. First, by sweeping *S* without re-tuning parameters, we reproduce the canonical sequence of dynamical regimes (self-sustained pulses, damped pulses, plateau), thereby confirming systems-level predictions and single-cell measurements on p53 oscillatory control (Bar-Or et al., 2000; Lahav et al., 2004; Ciliberto et al., 2005; Proctor & Gray, 2008; Geva-Zatorsky et al., 2006; Batchelor et al., 2008, 2011). Within the oscillatory domain, sustained p53 activity confirms the preferential engagement of *p21* and arrest/senescence programs (Harper et al., 1993; Xiong et al., 1993; Abbas & Dutta, 2009; Stewart-Ornstein & Lahav, 2017; Hanson et al., 2019). Outside this domain, the collapse of oscillations toward a steady state agrees with strong *PUMA* induction and an apoptotic bias (Nakano & Vousden, 2001; Jeffers et al., 2003; Kracikova et al., 2013; Zhang et al., 2007).

Second, our explicit (*S*, Σ) bifurcation map complements prior descriptions of the p53–Mdm2 axis by adding a tunable STAT1 dimension that shifts the Hopf band and reshapes fate partitions. In particular, increasing Σ both weakens *Mdm2*-mediated p53 removal and co-activates effector promoters, which aligns with reports that interferon/STAT signals feed back into p53 networks and IFN programs (Levy & Darnell, 2002; Brzostek-Racine et al., 2011; Feng et al., 2020). Our timing experiments further show that sustained or promptly rising Σ(*t*) accelerates early-gate crossing and increases transient PUMA load, while brief or delayed Σ(*t*) yields milder, often reversible effects. This extends observations on stimulus-dependent decoding of p53 dynamics (Batchelor et al., 2011; Purvis et al., 2012; Becker et al., 2010; Wang et al., 2023) by making the STAT1 time-profile an explicit gain/timing knob.

At population scale, injecting realistic heterogeneity into *S* and Σ reproduces lineage-specific fate biases without re-fitting biochemical parameters: HCT116-like populations concentrate on *p21* -dominated arrest/senescence, U2OS-like populations show enhanced *PUMA* responsiveness and apoptosis, and MCF-7-like populations remain mixed. This confirms and refines the idea that fate depends not only on p53 amplitude but also on target stability and feedback efficiency (Hanson et al., 2019; Stewart-Ornstein & Lahav, 2017; Sun et al., 2009). It also complements literature on S-phase damage and quiescence decisions via p21 (Barr et al., 2017), by embedding these outcomes in a two-input (*S*, Σ) landscape that predicts cohort-level splits.

Regarding STAT1, our results converge with evidence that prolonged STAT1 activity can potentiate cytotoxic and apoptotic programs (Heitmeier et al., 1999; Giles et al., 2017; Kalliara et al., 2022; Townsend et al., 2004) and are consistent with STAT1–p53 cooperativity on damage-induced apoptosis. At the same time, the literature also documents context-dependent, sometimes pro-tumoral STAT1 roles; our model adds that the temporal form of Σ(*t*) (sustained versus pulsed) is a key discriminator that may reconcile seemingly opposing observations (Meissl et al., 2017). Mechanistically, placing the STAT1 dimension alongside *S* clarifies how upstream cytokine cues reshape intracellular decoding by the p53 network and its effectors.

Finally, our framework suggests two natural extensions. A compartmental version will separate nuclear and cytosolic pools and translocation feedbacks, building on established transport models (Dimitrio, 2012; Eliaš et al., 2014). Coupling this stress–response module to a mechanistic JAK–STAT pathway model would close the loop from extracellular ligands to intracellular Σ(*t*) and fate decoding (Levy & Darnell, 2002), enabling predictions testable by combining controlled interferon waveforms with DNA-damage protocols and single-cell readouts of p53, p21, and PUMA.

## Supplementary Information

### S1. The model

#### S1.1. Calibration of the stress scale S

We sought a single, dimensionless stress variable *S* ∈ [0, 25] that captures how UV and *γ* irradiation stabilize *p*53 by weakening its Mdm2–mediated ubiquitination. Table S1 compiles *empirical anchors* from the literature: for representative UV or *γ* doses we list the measured *p*53 half–life *t*_1*/*2_, the corresponding effective ubiquitination rate *k*_*u*_ ≃ ln 2*/t*_1*/*2_, and the *S* value assigned to that condition. These anchors cover the full range from basal conditions (*S*=0) to severe damage (*S*∼25).

To complement these kinetic anchors, Table 1 in the main article summarizes the *phenotypic regimes* associated with increasing *S*: qualitative biological effects and the characteristic *p*53 dynamics observed near the corresponding *γ*-dose benchmarks. UV exposures were placed on the same *S* scale by matching *p*53 stabilization features (half–life/amplitude) to the nearest *γ*-dose regime.

**Table S1.**
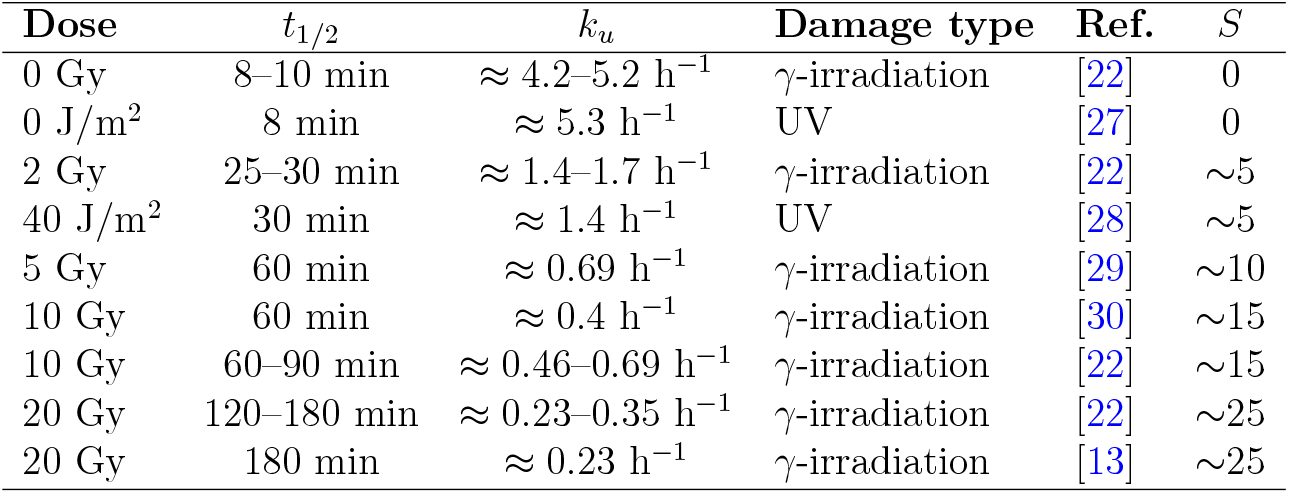
Mapping between dose, *p*53 half–life (*t*_1*/*2_), inferred ubiquitination rate (*k*_*u*_), and the assigned stress level *S*.

| Dose | $t_{1/2}$ | $k_u$ | Damage type | Ref. | $S$ |
| --- | --- | --- | --- | --- | --- |
| 0 Gy | 8–10 min | $\approx 4.2\text{--}5.2 \text{ h}^{-1}$ | γ-irradiation | [22] | 0 |
| 0 J/m<sup>2</sup> | 8 min | $\approx 5.3 \text{ h}^{-1}$ | UV | [27] | 0 |
| 2 Gy | 25–30 min | $\approx 1.4\text{--}1.7 \text{ h}^{-1}$ | γ-irradiation | [22] | $\sim 5$ |
| 40 J/m<sup>2</sup> | 30 min | $\approx 1.4 \text{ h}^{-1}$ | UV | [28] | $\sim 5$ |
| 5 Gy | 60 min | $\approx 0.69 \text{ h}^{-1}$ | γ-irradiation | [29] | $\sim 10$ |
| 10 Gy | 60 min | $\approx 0.4 \text{ h}^{-1}$ | γ-irradiation | [30] | $\sim 15$ |
| 10 Gy | 60–90 min | $\approx 0.46\text{--}0.69 \text{ h}^{-1}$ | γ-irradiation | [22] | $\sim 15$ |
| 20 Gy | 120–180 min | $\approx 0.23\text{--}0.35 \text{ h}^{-1}$ | γ-irradiation | [22] | $\sim 25$ |
| 20 Gy | 180 min | $\approx 0.23 \text{ h}^{-1}$ | γ-irradiation | [13] | $\sim 25$ |

#### Calibration procedure (summary)

In the model, stress modulates the effective ubiquitination activity through the factor

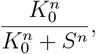

so that higher *S* reduces *p*53 degradation. We calibrated (*K*_0_, *n*) as follows:

1. Use Table S1 to anchor *S* values to measured *t*_1*/*2_ (and thus *k*_*u*_) under UV/*γ*.
2. For each anchor and for a grid of (*K*_0_, *n*), simulate the model and extract summary features of *p*53 dynamics (amplitude, period, damping) and stabilization.
3. Minimize the aggregate mismatch between simulated features and the regimes in Table 1 (e.g., oscillatory vs. plateau, strength of stabilization) while respecting the kinetic constraints from Table S1.
4. The best compromise was *K*_0_ = 12 and *n* = 6, which reproduces the transition from basal to pulsatile to stabilized *p*53 across the *S* range.

This procedure operationalizes the stress scale: the first table fixes kinetic anchors (half–life/ubiquitination), the second enforces dynamical phenotypes, and the fitted (*K*_0_, *n*) provide a consistent mapping from dose to *S* within the model.

### S2. Stability and bifurcation

#### S2.1. Existence, Uniqueness, and Positivity of Solutions

Existence, positivity, and uniqueness of solutions are given by the following theorem:

**Existence, uniqueness, and positivity of solutions**

Consider the system of differential equations defined on 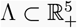:

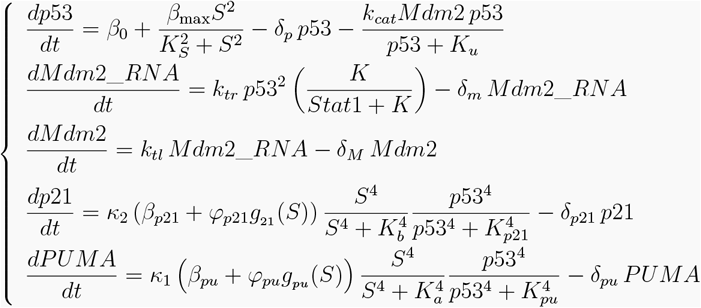

For any initial condition **x**_0_ ∈ Λ and any pair of continuous functions (*S*(*t*), Σ(*t*)), there exists a unique bounded solution **x**(*t*) defined on [0, +∞[, which remains in Λ for all *t* ≥ 0, i.e., all concentrations stay nonnegative at all times.

*Proof*. ***Existence of solutions***

All building blocks of *F* in **x** are 𝒞^∞^ on Λ (polynomials, products, and quotients whose denominators are of the form *p*53 + *K*_*u*_, *S*^4^ + *K*^4^, *p*53^4^ + *K*^4^, etc., with strictly positive constants). Hence, for each *t* ≥ 0 and continuous *S*, Σ, the map **x** → *F* (*t*, **x**) is 𝒞^1^ on Λ, thus locally Lipschitz. By the Picard–Lindelöf theorem, there is a unique local solution from any **x**_0_ ∈ Λ.

#### Positivity

We examine the vector field on the boundary of the non-negative orthant

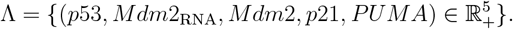

At the boundary *p*53 = 0, the *p*53 equation gives

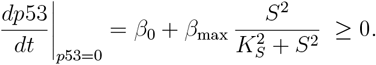

At the boundary *M dm*2_RNA_ = 0, the *M dm*2_RNA_ equation gives

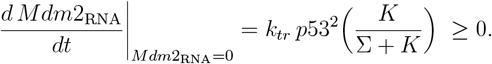

At the boundary *M dm*2 = 0, the *M dm*2 equation gives

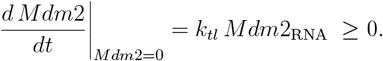

At the boundary *p*21 = 0, the *p*21 equation gives

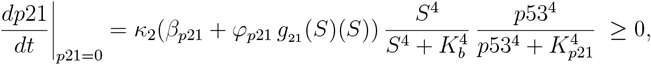

At the boundary *PUMA* = 0, the *PUMA* equation gives

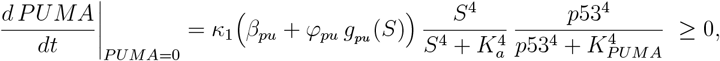

with *g*_*pu*_ (*S*) ≥ 0.

In each case the vector field points inward (or is tangent) on the coordinate hyperplanes. Therefore, Λ is a **positively invariant set** for the system.

### Boundaries for the core module. On *p*53 = 0

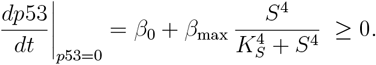

On *M dm*2_RNA_ = 0,

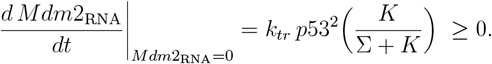

On *M dm*2 = 0,

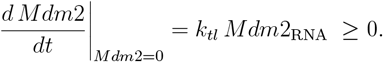

**Boundaries for the effectors**. On *p*21 = 0,

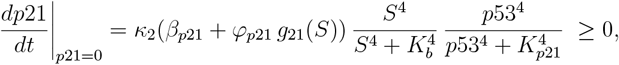

where *g*_21_(*S*) ≥ 0 is the non–negative Hill term used in the model.

On *PUMA* = 0,

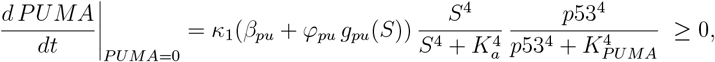

with *g*_*pu*_(*S*) ≥ 0 the stress–dependent Hill factor used for *PUMA*.

Since each boundary derivative is non–negative, trajectories starting in Λ remain in Λ for all forward times; i.e., Λ is a positive invariant set for the system.

#### Boundedness

We first recall (differential) Grönwall’s inequality, which we will use repeatedly.

*The Grönwall–Bellman inequality (differential form)*. Let *u*(*t*) be continuously differentiable on [0, *T* ] and satisfy

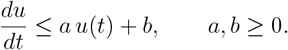

Then, for all *t* ∈ [0, *T* ],

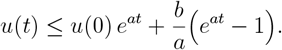

If *a* = 0, then *u*(*t*) ≤ *u*(0) + *b t*.

Below we use that every factor of the form 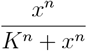 lies in [0,1], and that 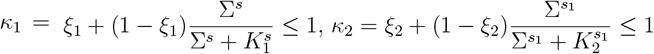.

i. *p*53(*t*) **is bounded**.

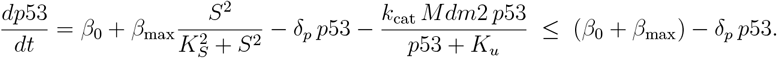

By Grönwall,

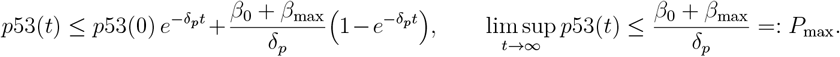
ii. *M dm*2_RNA_(*t*) **is bounded**.

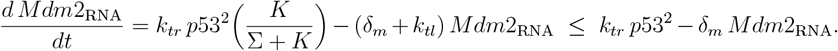

Using *p*53(*t*) ≤ *P*_max_ gives

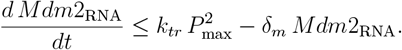

Thus,

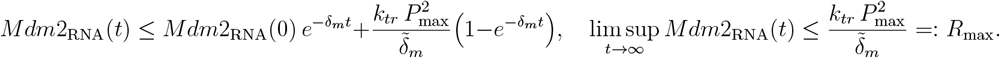
iii. *M dm*2(*t*) **is bounded**.

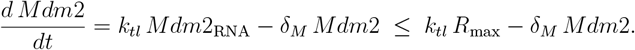

Therefore,

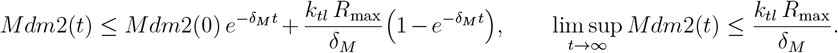
iv. *p*21(*t*) **is bounded**.

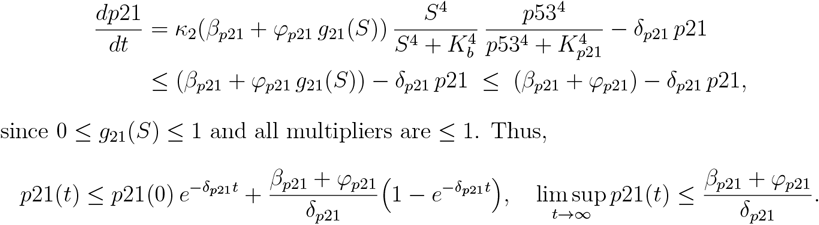

since 0 ≤ *g*_21_(*S*) ≤ 1 and all multipliers are ≤ 1. Thus,

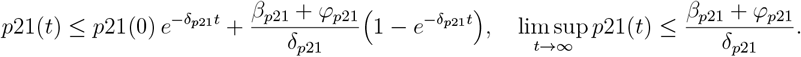
v. *PUMA*(*t*) **is bounded**.

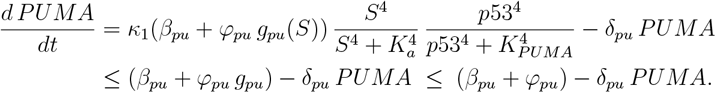

Hence,

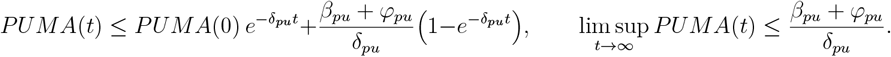

Combining (i)–(v), every component of the solution remains uniformly bounded for *t ≥* 0. Together with positivity (shown above), this implies that trajectories remain in a compact, positively invariant subset of the nonnegative orthant. Moreover, since the right-hand side is 𝒞^1^ in the state variables on Λ (hence locally Lipschitz), the Picard–Lindelöf theorem ensures existence and *uniqueness* of a local solution; the a priori bounds extend it to a *unique global* solution for any initial condition in Λ. This establishes that the problem is *well posed* both mathematically and biologically.

#### S2.2. Global sensitivity analysis (LHS–SRC, final–mean outputs)

##### Method

We quantified how model parameters shape the dynamics of the *p*53–Mdm2 core and of the effectors *p*21 and *PUMA* using *Standardized Regression Coefficients* (SRC) on samples generated by *Latin Hypercube Sampling* (LHS).

#### S2.3. Bifurcation loci in the (S, Σ) plane

Fig. S2 shows the real part of the leading eigenvalue as *S* is varied for several fixed Σ. A positive real part indicates an unstable fixed point and, here, the onset of a Hopf oscillatory regime. For Σ = 0 (blue) the Hopf window spans *S* ≈ 3.6–22.7. At Σ = 100 (red) it shrinks to *S* ≈ 4.3–15.1, and at Σ = 250 (green) to *S* ≈ 5.9–12.1. For Σ = 500 (orange) the curve remains negative for all *S* explored, hence no Hopf crossing. Two systematic trends emerge: (i) the onset 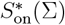 (dotted lines) moves to the right as Σ increases; (ii) the termination 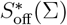 (dash-dotted lines) moves left. Consequently, the oscillatory band narrows with Σ.

**Figure S1.**
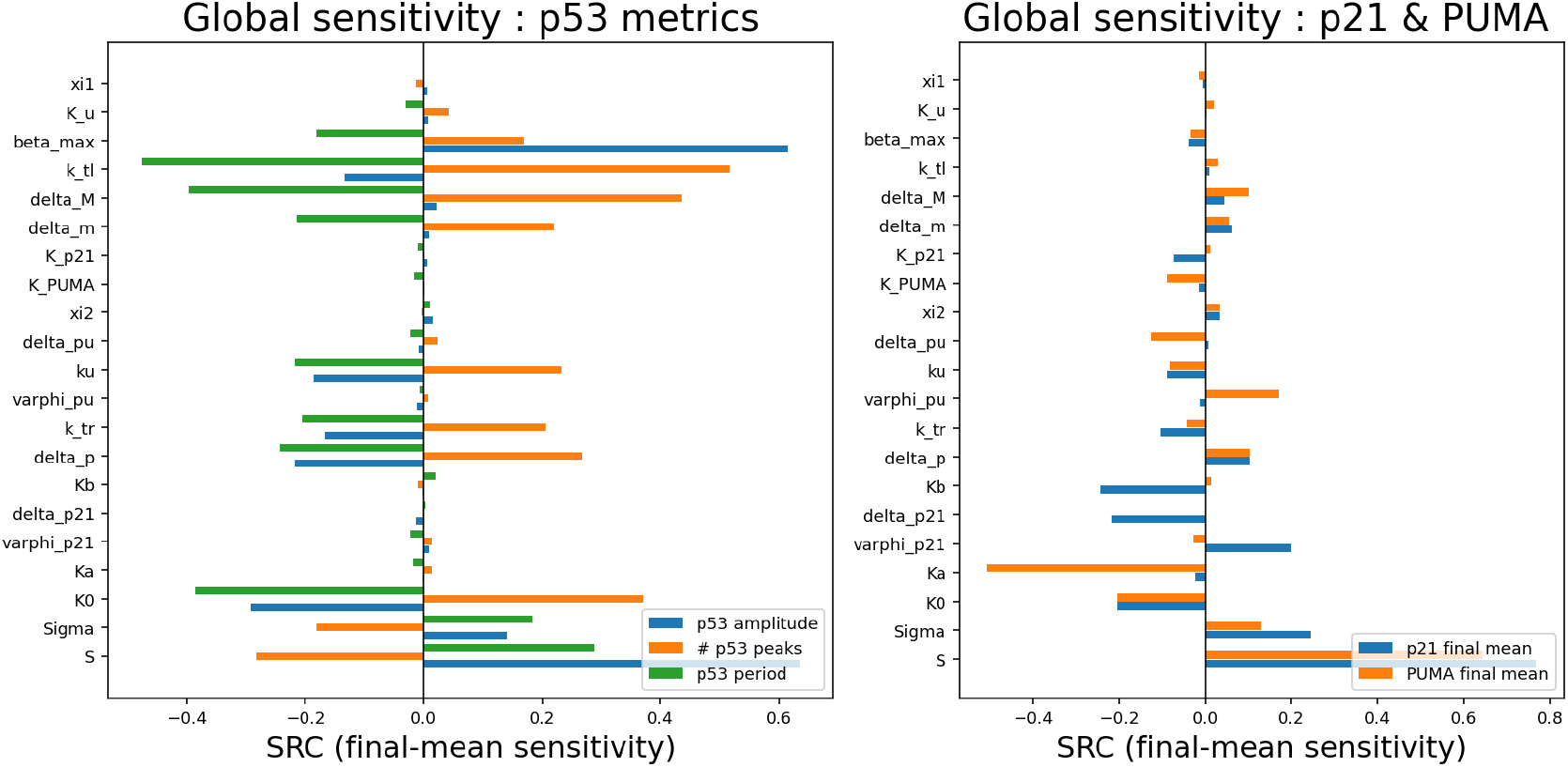
Global sensitivity by final-mean outputs (SRC). LHS (±30%) and SRC on (**left**) p53 metrics (amplitude, #peaks, period) and (**right**) final-mean p21 and PUMA (60–72 h). Positive bars indicate parameters that *increase* the output when raised; negative bars indicate the opposite. Results reveal partly distinct regulators for p53 oscillatory behaviour versus effector accumulation.

**Figure S2.**
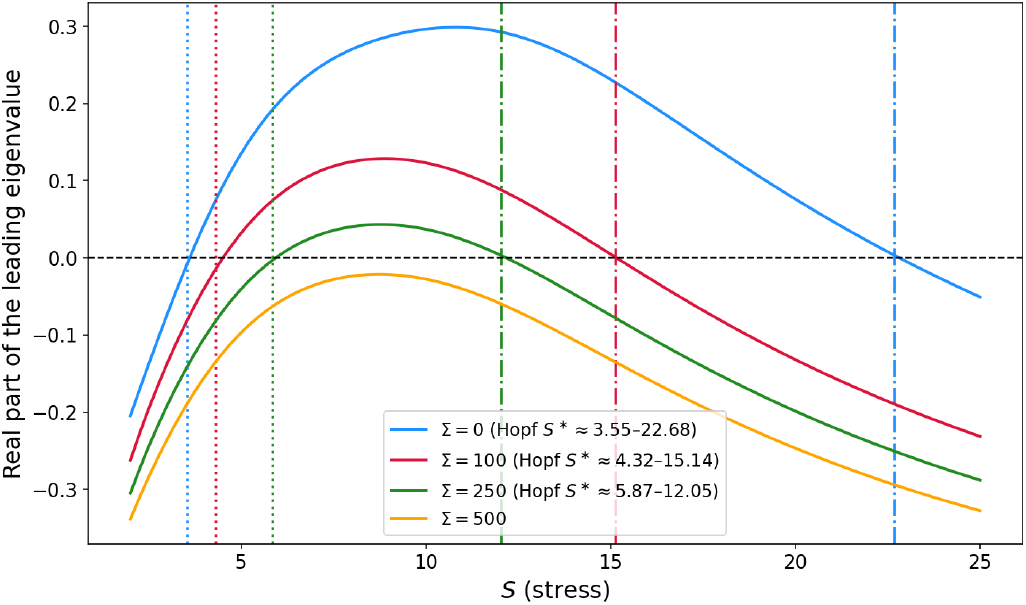
Influence of Σ on the Hopf threshold *S*^*\**^ in the p21 model. Real part of the dominant eigenpair at 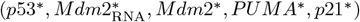 while scanning *S* for multiple Σ. Vertical dotted lines mark 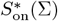; vertical dash-dotted lines mark 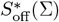.

**Table S2.** Most influential parameters from the SRC analysis (final–mean outputs). Arrows indicate the direction of the standardized regression coefficients (SRC). Only the dominant effects (largest |SRC| in Fig. S1) are listed.

| p53 dynamics (SRC on amplitude, #peaks, period) |  |
| --- | --- |
| Amplitude ↑ | $\beta_{\max} \uparrow, K_0 \uparrow, \Sigma \uparrow$ (strong positive SRCs): stronger production and a higher ubiquitination threshold raise *p53* pulse height; STAT1 ($\Sigma$) amplifies *p53* availability. |
| Amplitude ↓ | $\delta_p \uparrow$ (faster *p53* turnover) and faster Mdm2 turnover ($\delta_M, \delta_m \uparrow$) reduce pulse height by strengthening removal. |
| # Peaks ↑ | $\beta_{\max} \uparrow, k_{tl} \uparrow$ (more Mdm2 translation closes the negative loop faster): more frequent or more sustained oscillations. |
| # Peaks ↓ | $\delta_M, \delta_m \uparrow$ (weaker RNA/protein persistence in the Mdm2 arm) and very large $K_0$ beyond the oscillatory window. |
| Period ↑ | $K_0 \uparrow, \Sigma \uparrow$: a higher ubiquitination threshold and stronger STAT1 lengthen the cycle. |
| Period ↓ | $k_{tl} \uparrow$ and $\delta_p \uparrow$: a tighter/faster feedback shortens the inter-peak interval. |
| Effectors (SRC on final-mean over 60–72 h) |  |
| *p21* final-mean ↑ | $S \uparrow, \varphi_{p21} \uparrow, \Sigma \uparrow$: stress and STAT1 boost transcriptional drive; higher promoter gain increases accumulation. |
| *p21* final-mean ↓ | $\delta_{p21} \uparrow$ (faster decay), $K_b \uparrow$ (higher stress half-activation makes *p21* less sensitive at low $S$). |
| *PUMA* final-mean ↑ | $S \uparrow, \Sigma \uparrow, \varphi_{pu} \uparrow$: genotoxic stress and STAT1 coactivation promote robust *PUMA* expression. |
| *PUMA* final-mean ↓ | $\delta_{pu} \uparrow$ (faster decay), $K_a \uparrow$ (higher stress threshold; note $K_a > K_b$, so *PUMA* engages at higher $S$ than *p21*). |

These trends can be understood as follow: STAT1 enters the core loop through the factor 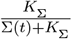 in the *M dm*2 equation; increasing Σ lowers this factor, reducing *Mdm*2 production and thus the effective *p*53 → *Mdm*2 negative-feedback gain. Sustained oscillations require enough loop gain and delay; when Σ is high the fixed point is more strongly damped, so a larger stress is needed to destabilize it (rightward shift of 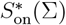). At the same time, in our model *S* decreases *k*_cat_ (the *M dm*2-mediated p53 removal rate), which weakens the feedback at high stress and brings the system back to stability—explaining the leftward shift of 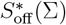 and the progressive collapse of the oscillatory window.

#### Interpretation

*Left: p*53 amplitude (60–72 h) forms a Hopf “tongue” at intermediate *S*. As Σ increases, this oscillatory band *shifts* to higher *S* and *narrows*, ultimately extinguishing oscillations at large Σ. *Middle: p*21 amplitude (60–72 h) mirrors the *p*53 tongue but is narrower and decays faster with Σ; when Σ stabilizes *p*53 on a high plateau, the mean of *p*21 can remain moderate/high while its amplitude is low. *Right:* max(PUMA) (0–100 h) increases monotonically with *S* and Σ, reflecting stress sensitization and STAT1 co-activation. *Fate reading:* the oscillatory band aligns with arrest (ARR); its progressive closure with growing Σ promotes damped/intermediate responses (consistent with SEN) and, at high *S* and/or Σ, large PUMA peaks bias decisions toward apoptosis (APO).

#### Interpretation

Figure S4 shows that, in addition to the stress input *S*, both the Mdm2 degradation rate *δ*_*M*_ and the saturation constant *K*_0_ act as bifurcation parameters for the p53 network. For small values of *δ*_*M*_, oscillatory regimes can emerge, and their extent depends moderately on the level of STAT1 (Σ). Similarly, low *K*_0_ values may also trigger a Hopf bifurcation, but again the influence of Σ is weaker than in the case of stress *S*. Overall, these results indicate that while *K*_0_ and *δ*_*M*_ shape the boundaries of oscillatory domains, STAT1 mainly fine-tunes them, and the dominant bifurcation control remains associated with the stress level *S*.

**Figure S3.**
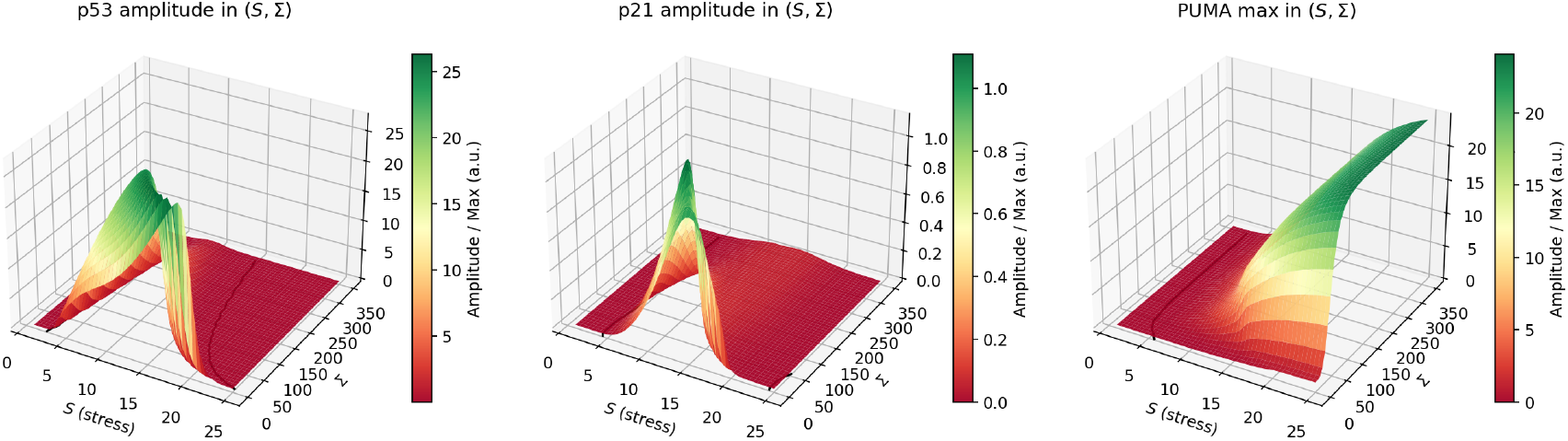
Hopf bifurcation for *K*_0_ and *δ*_*M*_ .

**Figure S4.**
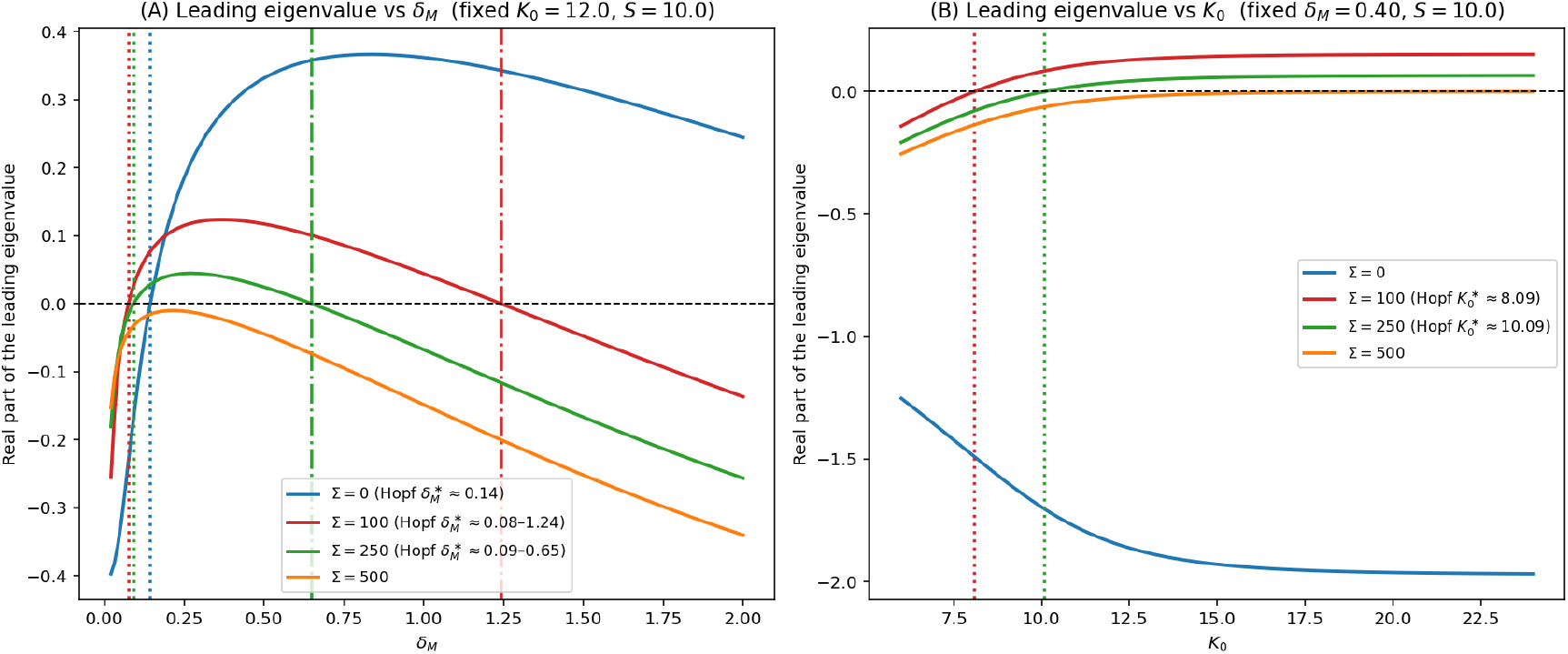
Hopf bifurcation for *K*_0_ and *δ*_*M*_ .

### S3. Fate criterion, Calibration details and evaluation metrics

#### Calibration of stress-agnostic stationary-index cutoffs α_f_

We fix a single pair *α*_*f*_ = (*α*_*f*1_, *α*_*f*2_), shared by all stress levels *S*.

A. **Stationary window and low thresholds** Define the stationary window 𝒲 = [60, 72] h and the time averages

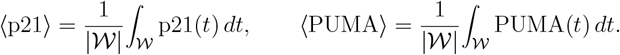

We use three global thresholds: *µ*_21_ (low p21), *µ*_PU_ (low PUMA), and *µ*_*S*_ *> µ*_21_ (senescence-level p21). The stationary index *I*_*f*_ is evaluated only when both species exceed their low thresholds.
B. **Global upper bounds for stationary concentrations** To remain stress-agnostic, we estimate 95th-percentile upper envelopes across stresses:

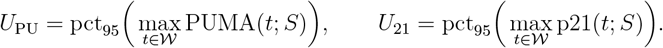
C. **Admissible interval for the stationary index** When both species maintain high stationary levels, we define the balance index

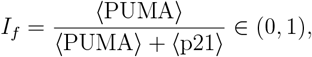 which measures the relative contribution of pro-apoptotic PUMA compared to the total (PUMA + p21) pool. Let us denote *A* = ⟨PUMA⟩ and *B* = ⟨p21⟩. Under the admissible ranges

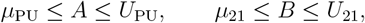

we seek the interval of all possible values of *I*_*f*_ (*A, B*) = *A/*(*A* + *B*). Since

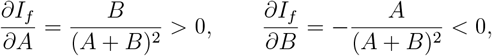

the function *I*_*f*_ (*A, B*) is strictly *increasing* in *A* and strictly *decreasing* in *B*. Therefore, the extremal values of *I*_*f*_ over the admissible rectangle [*µ*_PU_, *U*_PU_] × [*µ*_21_, *U*_21_] occur at opposite corners:

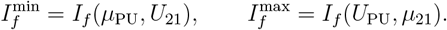

Hence,

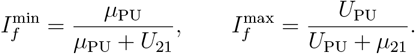

By construction, 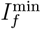 corresponds to the most p21-dominant configuration (low PUMA, high p21), while 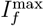 corresponds to the most PUMA-dominant configuration (high PUMA, low p21). The admissible interval 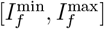 thus bounds all biologically plausible stationary balances between pro-apoptotic and cytostatic dominance, and provides a stress-agnostic reference for the calibration of the global cutoffs *α*_*f*1_, *α*_*f*2_.

(D) Choosing the two cutoffs *α*_*f*1_, *α*_*f*2_ Two options are considered:

- **Heuristic (fast)**. With a safety margin *δ* ∈ [0.02, 0.05],

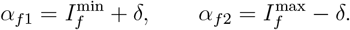

This choice partitions the admissible interval into three regions: *I*_*f*_ *< α*_*f*1_, *α*_*f*1_ ≤ *I*_*f*_ *< α*_*f*2_, and *I*_*f*_ *≥ α*_*f*2_, which correspond respectively to cycle/viability, arrest, and apoptotic regimes.
- **Minimax (robust)**. Perform a grid search over 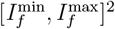 with *α*_*f*1_ *< α*_*f*2_, minimizing the worst per-stress classification error:

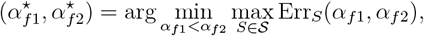

where Err_*S*_ = 1 − F1 or 1 − accuracy on the population at stress *S*.

**(E)Evaluation metrics (precision, recall, F1)** For each class (one-vs-rest),

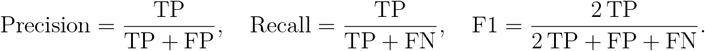

For multi-class fate (CY/ARR/APO), we report macro- and weighted-F1 scores; micro-F1 equals overall accuracy.

**(F) Mapping from** *I*_*f*_ **to fate (explicit rule)** With fixed global cutoffs 0 *< α*_*f*1_ *< α*_*f*2_ *<* 1,

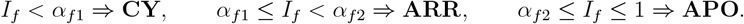

*Note:* senescence is now encompassed within the arrest band when *p*21 exceeds *µ*_*S*_ but *I*_*f*_ *< α*_*f*2_, reflecting cytostatic rather than apoptotic dominance.

**(G) Validation (per stress)**. For each stress level, compute per-class F1 scores; accept the global pair *α*_*f*_ if all strata exceed a floor (e.g. F1 ≥ 0.70) and the across-stress variation remains moderate (range ≤ 0.10). Leave-one-stress-out validation avoids leakage.

*Early index cutoffs α*_*p*_ *(analogous procedure)*. For the early overlap window *J* ^*⋆*^,

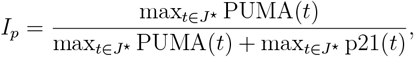

and the same two-cutoff strategy (*α*_*p*1_, *α*_*p*2_) is applied to delineate early-cycle vs early-apoptotic dominance. Validation mirrors the stationary case. Our cell fate criterion reads:

##### Supplementary Box 1

Cell-fate decision criterion (two global cutoffs)

**Setup**. Fix a single set of thresholds, independent of stress *S*:

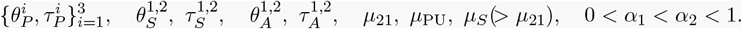

Let *J*(*τ* ) be any contiguous window of length ≥ *τ* .

**(1) Early decision (before 60 h)**.

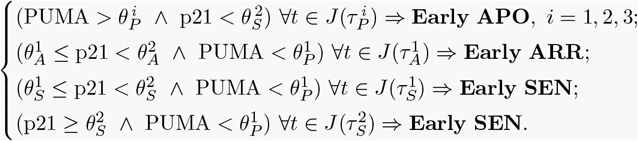

**(2) Early index (PUMA–p21 overlap)**. If there exists 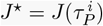 with 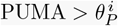 and 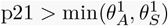 for all *t* ∈ *J*, define

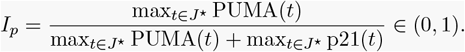

Then classify with global cutoffs 0 *< α*_1_ *< α*_2_ *<* 1:

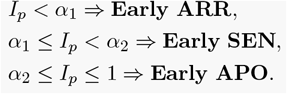

**(3) Stationary decision (60–72 h)**. With ⟨·⟩ = ⟨·⟩_[60,72]_:

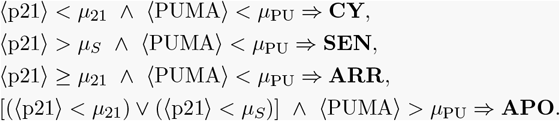

**(4) Stationary index (both high)**. If (⟨p21⟩ ⩾ *µ*_21_) ∨ (⟨p21⟩ ⩾ *µ*_*S*_) and ⟨PUMA⟩ ⩾ *µ*_PU_, define

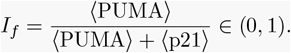

Then classify with the same global cutoffs:

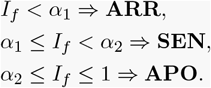

**(5) Effective decision (severity-first)**. Let Early ∈ {None, ARR, SEN, APO} and Stationary ∈ {CY, ARR, SEN, APO}. Return the effective fate

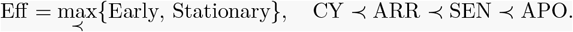

### S4. Simulations

Here are simulations made on generic single cells in addition to the ones in the main article. The following table gives the threshold values used in the single generic cell simulations.

**Table S3.** Thresholds and durations used in the cell-fate decision criterion, with both the *global* stationary-index cutoffs (*α*_*f*1_, *α*_*f*2_) = (0.24, 0.50) and the early-index cutoffs (*α*_*f*1_, *α*_*f*2_) = (0.22, 0.50) obtained via the percentile-bounded calibration described in the text. Throughout, gates are stress-invariant: all cutoffs and early windows are fixed per archetype and reused for every stress level *S*, without per-*S* retuning.

| Name | Meaning (criterion scope) | Value |
| --- | --- | --- |
| **Global (stress-agnostic) index cutoffs** |  |  |
| $(\alpha_{p1}, \alpha_{p2})$ | Cutoffs for the *early* index $I_p$ (PUMA-anchored overlap); obtained via the percentile-bounded early calibration on $J^\star$ | (0.22, 0.50) |
| $(\alpha_{f1}, \alpha_{f2})$ | Cutoffs for the *stationary* index $I_f$ (unique for all $S$); obtained via the percentile-bounded late calibration | (0.24, 0.50) |
| **Early thresholds (pre-60 h)** |  |  |
| $\theta_P^{1,2,3}$ | PUMA thresholds (early apoptosis) | (10, 12, 14) nM |
| $\tau_P^{1,2,3}$ | Minimal required durations for PUMA (APO) | (6, 4, 2) h |
| $\theta_A^{1,2}$ | $p21$ thresholds (early arrest) | (6, 8) nM |
| $\tau_A^{1,2}$ | Minimal required durations (arrest) | (6, 4) h |
| $\theta_S^{1,2}$ | $p21$ thresholds (early senescence) | (10, 13) nM |
| $\tau_S^{1,2}$ | Minimal required durations (senescence) | (6, 4) h |
| **Stationary gates (60–72 h)** |  |  |
| $\mu_{21}$ | Minimal effective $p21$ level | 2 nM |
| $\mu_{PU}$ | Minimal effective PUMA level | 4 nM |
| $\mu_S$ | Robust senescence threshold (final) | 14 nM |
| **Time windows** |  |  |
| **Early window** | Interval used to test early engagement | $t < 60$ h |
| **Stationary window** | Averaging interval for stationary decision | [60, 72] h |

#### S4.1. Cell fate under constant stress and constant STAT1

We here consider constant levels of both stress and Stat1

**Figure S5.**
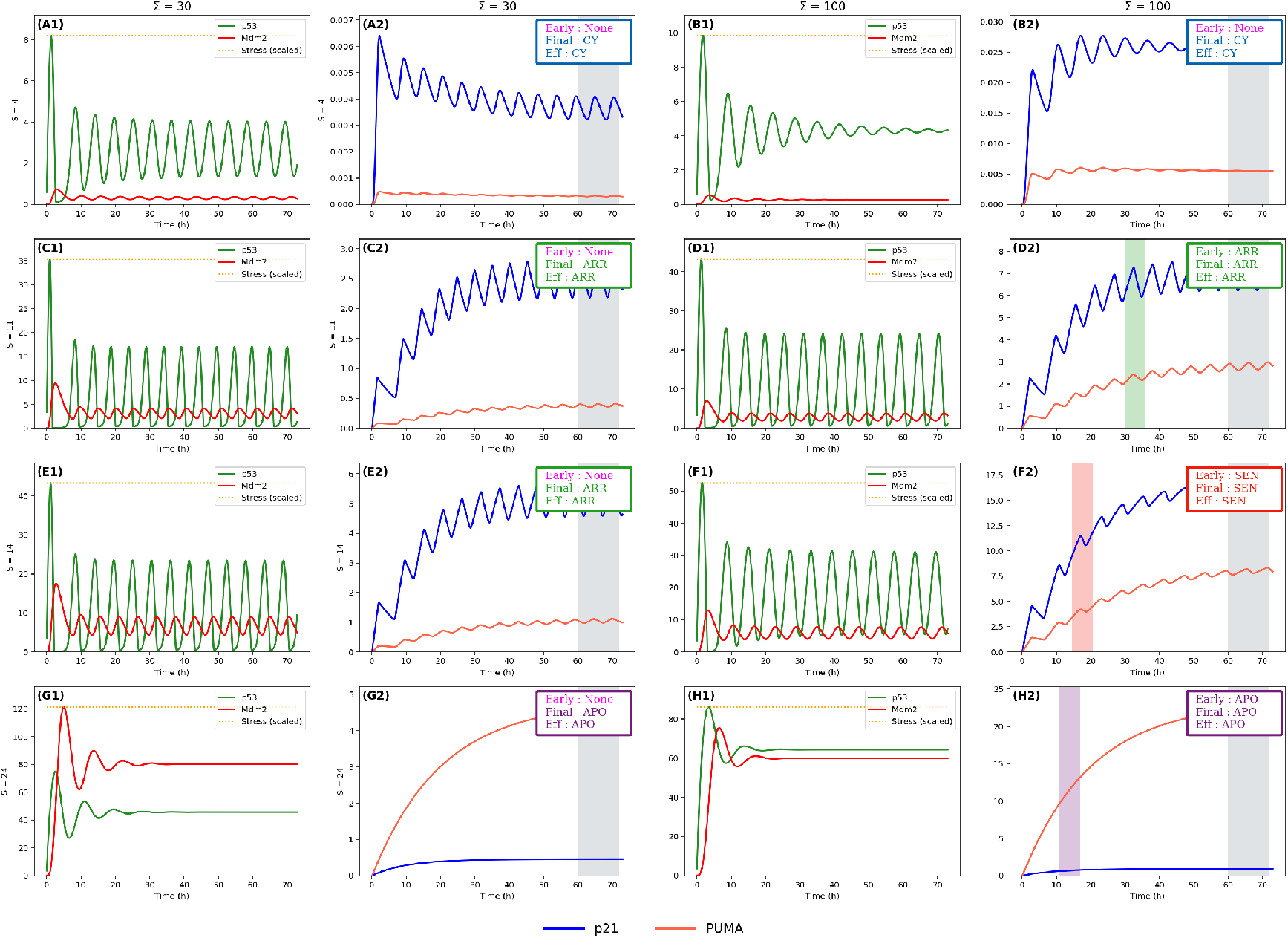
Time courses of *p*21 and PUMA for different fixed STAT1 levels Σ. Horizontal lines: thresholds; grey band: 60–72 h; tags indicate the effective fate.

*Interpretation*. With constant inputs *S* and Σ, trajectories relax to a fixed attractor—no late reversals occur. When a coloured vertical band appears on the PUMA/*p*21 panel, it marks the time at which the *early* rule is first satisfied; under constant inputs this early label always matches the stationary label (60–72 h). If no band appears, fate is read from the stationary window only. At fixed *S*, raising Σ (STAT1) accelerates PUMA/*p*21 production and weakens the *p*53–Mdm2 negative feedback, so commitment (when present) tends to occur earlier and oscillations are more strongly damped; in some cases the attractor class (oscillatory vs. plateau) changes and the fate switches.

*S* = 4 (low stress): The *p*53–Mdm2 loop decays to a low fixed point. Both *p*21 and PUMA remain well below threshold; no early band is detected. The cell keeps cycling for Σ = 30 and for Σ = 100 (A2,B2: CY).

*S* = 11 (moderate stress): The core shows a stable pulse train (limit cycle). For Σ = 30, *p*21 crosses the arrest gate while PUMA stays sub-apoptotic—*arrest* without an early band (C2: ARR). For Σ = 100, the same outcome is reached earlier and is flagged by an early green band (D2: ARR→ARR).

*S* = 14 (higher stress): With Σ = 30 the system remains pulsatile and *p*21 settles in the arrest range—*arrest* with no early band (E2: ARR). With Σ = 100 the stronger STAT1 drive damps the oscillations to an intermediate plateau and maintains higher *p*21, producing *senescence* with an early red band (F2: SEN→SEN).

*S* = 24 (extreme stress): Trajectories collapse rapidly to a high plateau with strong PUMA and relatively low *p*21. Both conditions end in *apoptosis*; however, only the higher STAT1 produces an early apoptotic band (G2: APO without early band; H2: APO with an early purple band).

Overall, constant-input experiments confirm that fate is governed by the steady attractor reached under (*S*, Σ), while Σ primarily modulates the timing of commitment and, when it reshapes the attractor (oscillations → plateau), can shift ARR↔SEN.

#### S4.2. Cell fate under time-varying stress

We here assume stress is a time-dependant function exhibiting different profiles.

**Figure S6.**
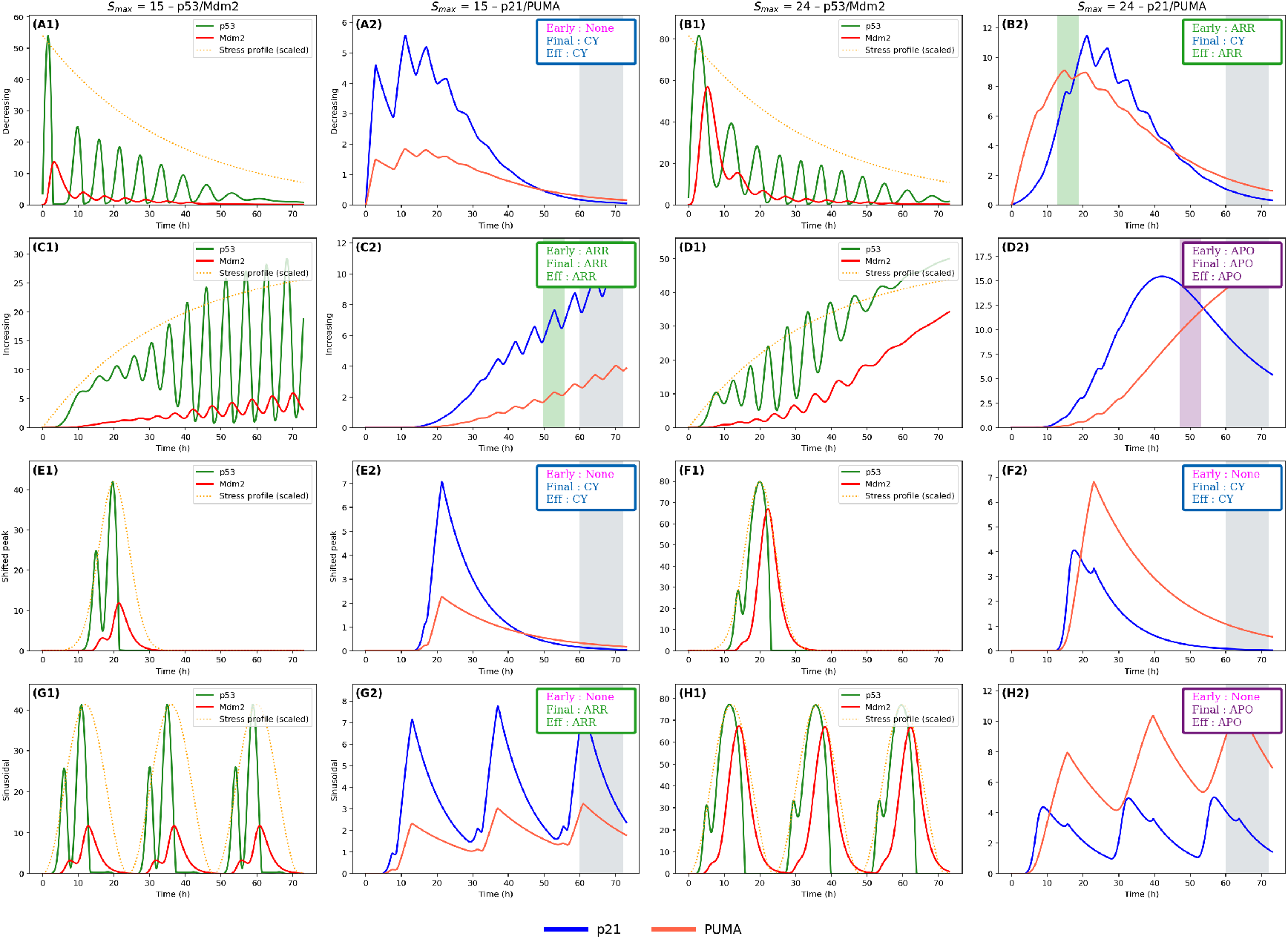
Time courses for four stress profiles *S*(*t*) (decreasing, increasing, peak, sinusoidal) at *S*_max_=15 and 24.

#### Interpretation (time–varying stress, Σ = 100)

Coloured vertical bands mark the first time the *early* rule is satisfied; when no band appears, fate is decided only from the stationary window (60–72 h). We report fates as continued cycling (CY), cell–cycle arrest (ARR), senescence (SEN), and apoptosis (APO).

#### Decreasing exponential

A rapidly decaying input is too brief to sustain effectors. At *S*_max_ = 15 there is no early band and the stationary decision is CY (A2). At *S*_max_ = 24 a short *p*21 overshoot triggers an early ARR band, but the signal vanishes before 60–72 h, so the stationary decision is CY (B2).

#### Increasing ramp

A steadily rising input prolongs activation. At *S*_max_ = 15, *p*21 crosses the arrest gate and the trajectory is ARR both early and at stationarity (C2). At *S*_max_ = 24, the ramp drives PUMA across its threshold; APO is reached early and remains APO in the stationary window (D2).

#### Shifted peak (narrow pulse)

A short stress pulse produces isolated spikes in the *p*53–Mdm2 loop that relax quickly. At both *S*_max_ = 15 and *S*_max_ = 24 there is no early band and the stationary decision is CY (E2, F2).

#### Sinusoidal waveform

Repeated input windows maintain *p*53*/p*21 pulsatility. At *S*_max_ = 15 there is no early commitment; the stationary window classifies ARR (G2). At *S*_max_ = 24 commitment often requires accumulation over several cycles—there may be no early band—yet PUMA integrates to a high plateau and the stationary decision is APO (H2).

Overall, waveform shape is decisive: sustained or rising inputs promote commitment (ARR at moderate intensity, APO at high intensity), brief/decaying inputs fail to meet stationary thresholds, and periodic inputs gate commitment in discrete windows, which explains why early and stationary calls can differ under the same peak stress.

#### S4.3. Sensitivity to initial p21 and PUMA

We wanted to see whether initial conditions do have any impact on cell fate.

**Figure S7.**
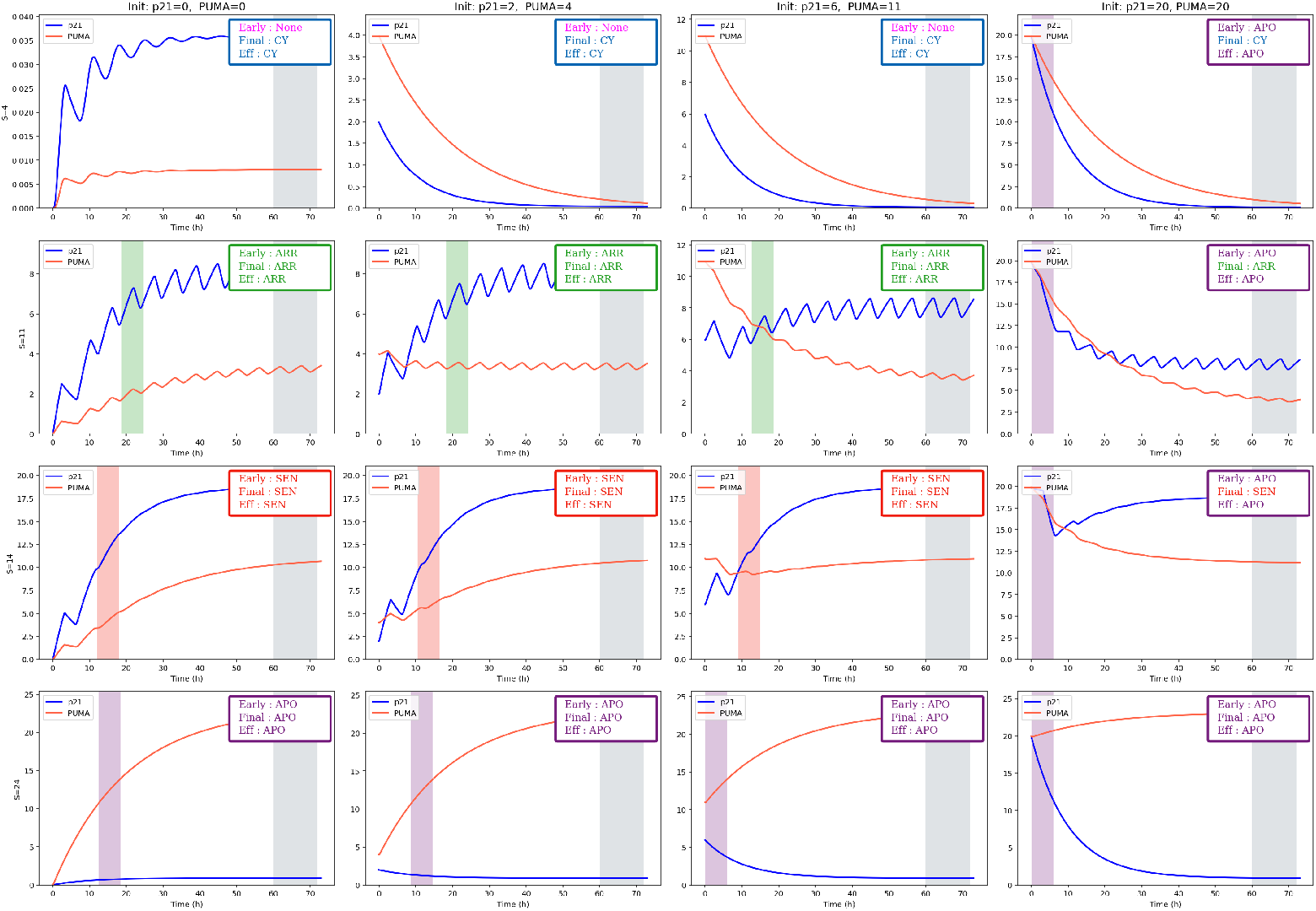
Effect of initial conditions for *p*21(0) and PUMA(0) on fate.

#### Interpretation

The grid varies the stress level across rows (*S* = 4, 11, 14, 24; Σ = 100 fixed) and the initial loads of *p*21/PUMA across columns (the last column is a deliberately extreme “preload”). Coloured vertical bands indicate the first time an early gate is crossed; when absent, fate is decided from the stationary window (60–72 h). For physiological initial conditions (first three columns), the effective fate is essentially set by *S*; initial conditions mainly shift the *timing* of commitment.

*S* = 4. Trajectories relax to a low attractor and no early gate fires in the physiological columns; the stationary decision is CY. Only the extreme PUMA preload triggers an early APO band that overrides the stationary CY by the severity priority, illustrating the rule rather than genuine multistability.

*S* = 11. The core remains on a sustained pulse train. *p*21 crosses the arrest gate around 30–45 h (green bands) and the stationary window confirms ARR in all physiological columns; initial loads mostly advance or delay the gate but do not change the fate.

*S* = 14. Pulses damp toward a mid–level plateau. In the physiological columns, *p*21 crosses the SEN gate (red band, ∼15–25 h) and the stationary call is also SEN. Under the extreme preload, PUMA briefly exceeds its early apoptotic gate, yielding an APO early band, but the stationary window classifies SEN; the effective fate is APO by the priority rule.

*S* = 24. Dynamics collapse to a high plateau with strong PUMA. An APO gate fires within 10–20 h in every column and the stationary decision is APO. Initial conditions only modulate the latency of commitment.

Overall, constant Σ with varied *S* reveals convergence to a unique late attractor at each dose (low oscillations → CY; sustained limit cycle → ARR; mid plateau → SEN; high plateau → APO). Initial conditions alter when thresholds are crossed but not which fate is ultimately reached, except under unrealistically large preloads where the early priority applies.

### S5. From single-cell rules to population outcomes

Our procedure to build a cell-line *archetype* follows a common protocol:

1. **Phenotypic targets (literature)**. For each representative *S*, we fix the expected Cycle/ARR/SEN/APO trends (window ∼48–72 h) and, when available, *early* signatures.
2. **Decision logic**. We adopt the criterion (early windows, stationary gates, and indices *I*_*p*_*/I*_*f*_ ) as defined in the main article Box **??**; the *θ/τ* and *µ*_•_ thresholds, as well as the *α* cutoffs, are deterministic and global.
3. **Archetype constants/nudges**. Certain amplitudes are adjusted *fixedly* to reflect lineage traits (e.g., MCF-7: PUMA dampening and slight *p*21 boost; U2OS: mild PUMA dampening). These *nudges* define the *central value* around which cells are sampled.
4. **Intra-population variability**. The parameters listed in the table below are drawn *independently* per cell from log-normal laws with the stated center and coefficient of variation (CV). We include a multiplicative noise factor on *S* (micro-variability), a draw for Σ (STAT1), and initial conditions IC (*p*21_0_, PUMA_0_).
5. **Calibration**. We tune first the *stationary gates* (*µ*_pu_, *µ*_21_, *µ*_*S*_), then the *early windows* (*θ, τ* ), and finally the *indices* (*α*_*p*•_, *α*_*f*•_). *Nudges* are only revised as a last resort.

#### S5.1. Archetypes

##### Computation

Given *N* single-cell simulations or trajectories *c* = {1, …, *N*}, we assign an effective fate Eff_*c*_ ∈ {CY, ARR, SEN, APO} per cell using our fate criterion, then report fractions

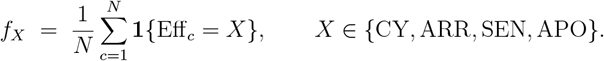

Uncertainty can be summarized by bootstrap CIs or by repeating with different random seeds.

## Supplementary Methods

*Validation against literature* intervals.

### Setup

For each stress level *S*, the simulation (SIM) outputs class proportions *p* = (*p*_CY_, *p*_ARR_, *p*_SEN_, *p*_APO_) Δ_3_ ∈ (Σ_*i*_ *p*_*i*_ = 1). The literature (LIT) provides *interval targets* ℛ = {*q* ∈ Δ_3_ : *L*_*i*_ ≤ *q*_*i*_ ≤ *U*_*i*_ for *i* = 1, …, 4} with 0 ≤ *L*_*i*_ ≤ *U*_*i*_ ≤ 1 and Σ_*i*_ *L*_*i*_ ≤ 1 ≤ Σ_*i*_ *U*_*i*_

1. *Feasibility and margin*. Define the per-class violation

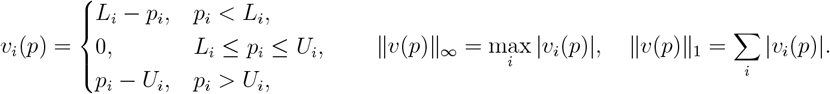

Feasible if and only if *v*(*p*) ≡ 0. A normalized margin (unitless) highlights which class is tightest:

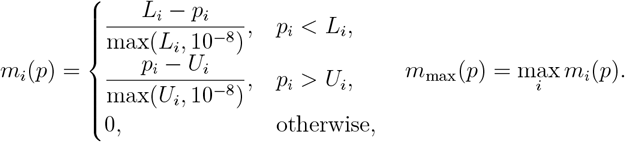
2. *Distance to the interval set (best achievable agreement)*. We measure the *closest* agreement to any vector within the LIT ranges:

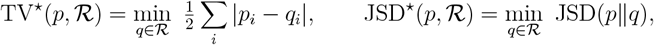

with JSD the Jensen–Shannon distance (base 2). Both minimizations are over a convex polytope (Δ_3_ intersected with the box [*L, U* ]); they can be computed by a small quadratic program (for JSD, using its convex proxy) or by projecting *p* onto ℛ in *ℓ*_1_ (for TV^*⋆*^).
3. *Minimal reallocation mass*. TV^*⋆*^ has a direct interpretation: it is the smallest fraction of cells that must be re-labeled (moved between classes) to bring SIM inside all LIT intervals. We also report the *edit plan* (which classes donate/receive).
4. *Sample-size aware coverage*. With *n* simulated cells, let 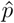 be the observed proportions and draw *B* bootstrap replicates 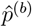 (multinomial). Define the interval *coverage*

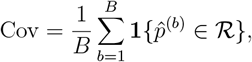

and per-class coverages 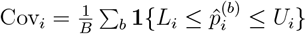. Coverage distinguishes tight agreement from incidental hits at small *n*.
5. *Class-wise standardized gap (sample-size free)*. To summarize how far the worst class sits from its admissible band,

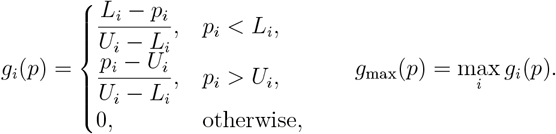

Here *g*_max_ = 1 means being one full band-width away from feasibility. *Decision rule (single stress level). Acceptable* if

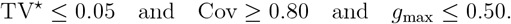

*Needs improvement* if TV^*⋆*^ *>* 0.05 or *g*_max_ *>* 0.50 but Cov [0.50, 0.80). *Not acceptable* otherwise. (Thresholds chosen for interpretability; they can be tuned in sensitivity analyses.)

### Aggregating across stresses

Report the above metrics per *S* and summarize with medians (and IQR) over *S*. For a single composite score, use the maximum distance to feasibility: max_*S*_ TV^*⋆*^(*p*(*S*), ℛ (*S*)), alongside the minimum coverage min_*S*_ Cov(*S*).

### Notes on implementation

We solve TV^*⋆*^ as a linear program on *q* with constraints *L*_*i*_ ≤ *q*_*i*_ ≤ *U*_*i*_, Σ_*i*_ ≥ *q*_*i*_ = 1, *q*_*i*_ ≥ 0. JSD^*⋆*^ is computed by a convex solver (or by grid search, given the low dimension). Bootstrap uses *B* = 200 by default; results are stable for *B* ≥ 100.

### S6. Simulations

#### S6.1. HCT116 archetype

Apoptosis and senescence already emerge at moderate stress.

**Figure S8.**
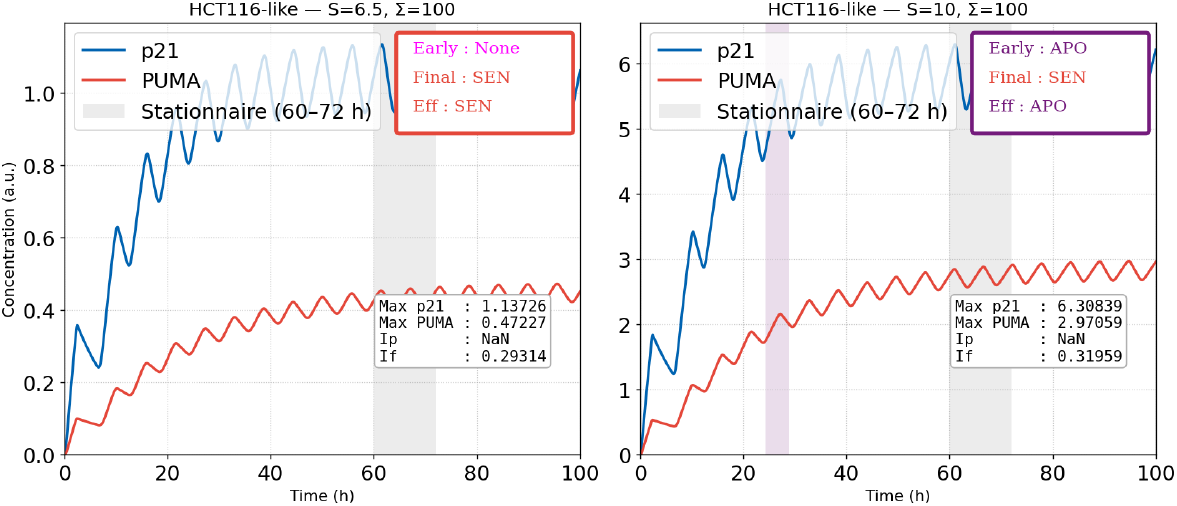
HCT116-like single–cell dynamics at two constant stresses (Σ = 100). **Left:** *S* = 6.5. p21 (blue) and PUMA (red) trajectories with the stationary window shaded (60™72 h). No stable early band is detected (*Early: None*); the late averages satisfy ⟨p21⟩ *> µ*_21_ and ⟨PUMA⟩ *< µ*_PU_, yielding **Final: SEN. Right:** *S* = 10. A transient PUMA-dominant early band is detected (**Early: APO**), while the late window again satisfies the senescence gate (**Final: SEN**). The severity-first rule therefore reports **Eff: APO**. Insets list peak concentrations and the stationary index *I*_*f*_ = ⟨PUMA⟩ */*(⟨PUMA⟩ + ⟨p21⟩); the shaded rectangles mark the decision windows (purple for the detected early band, grey for the stationary window).

**Figure S9.**
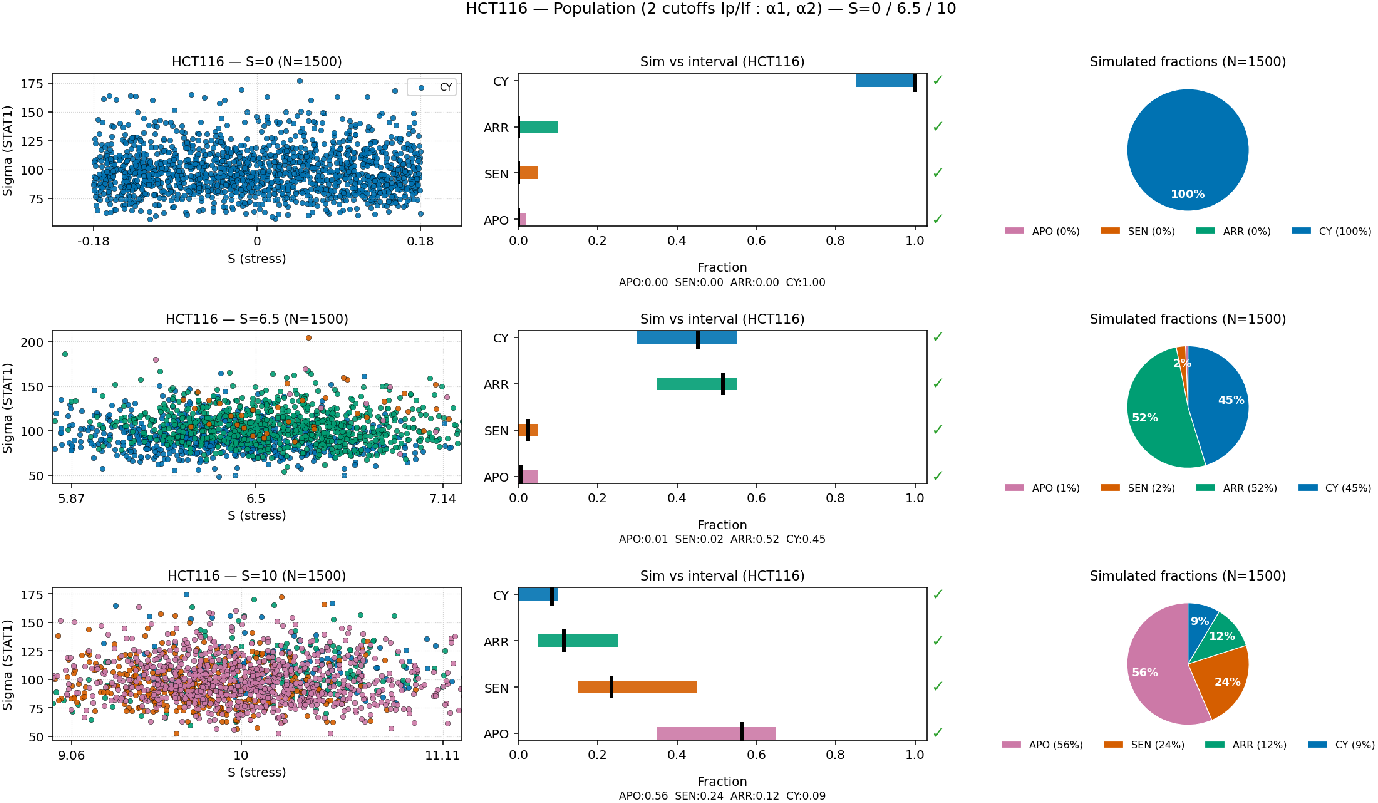
Population-level simulations showing the distribution of fates for the HCT116 archetype. The set of thresholds used for the simulations are given in Box**??**.

##### Supplementary Box 2

**HCT116-like** — Decision thresholds (global)

HCT116-decision-critere

**Early PUMA (APO):**

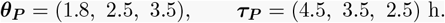

**Stationary gates (mean on** [60, 72] **h):**

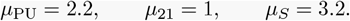

**Index cutoffs (used for** *I*_*p*_ **and** *I*_*f*_ ):

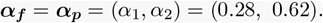

**Early** *p*21 **bands:**

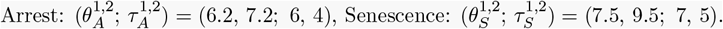

### Validation of population fractions against literature intervals (HCT116-like)

For each stress level *S* and fate *k* ∈ {CY, ARR, SEN, APO} the literature provides an admissible interval [*ℓ*_*S,k*_, *u*_*S,k*_]. From *N* simulated cells we estimate the fraction 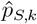 and evaluate:

i. *Interval coverage and normalized gap*.

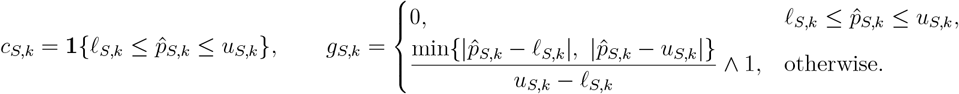

Aggregate per stress:

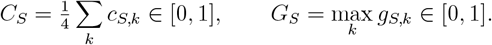
ii. *Uncertainty-aware checks*. For each (*S, k*) we compute a binomial Wilson 95% CI [lo_*S,k*_, hi_*S,k*_] for 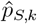 and record

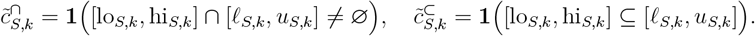
iii. *Decision rule (per stress)*.

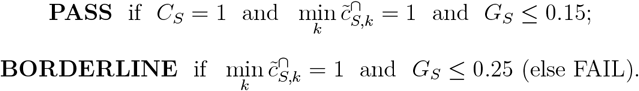

(Thresholds 0.15*/*0.25 are conservative and can be tuned.)

## Results (HCT116-like; N =1500 per S)

Across the three stress levels (*S*=0, 6.5, 10), the validation remains globally consistent and all cases are classified as **PASS**. At *S*=0, the population is entirely **cycling** (**CY** = 1.00), which lies well within the literature range and confirms the absence of premature activation under basal stress. At *S*=6.5, the fractions **ARR** ≈ **0.53, SEN** ≈ **0.01**, and **CY** ≈ **0.45** all fall within their admissible intervals, with **APO** remaining negligible—consistent with a dominant arrest response at intermediate stress. At *S*=10, the mixture shifts toward apoptotic and senescent outcomes (**APO** ≈ **0.55, SEN** ≈ **0.24, ARR** ≈ **0.12, CY** ≈ **0.09**), still inside the expected bands, reproducing the experimentally observed stress-induced transition from arrest to apoptosis. In all panels, simulated ticks (black bars) fall within or touch the colored literature intervals, and every class shows a green checkmark—indicating full interval coverage (*C*_*S*_=1.00), intersecting Wilson 95% confidence intervals, and small worst normalized gaps (*G*_*S*_ ≤ 0.10).

### S6.2. U2OS archetype

For this archetype, we aim to observe *senescence* and *apoptosis* already at low stress. We use a standard U2OS-like single cell; the biochemical model below (parameters and initial conditions) is fixed across simulations.

**Figure S10.**
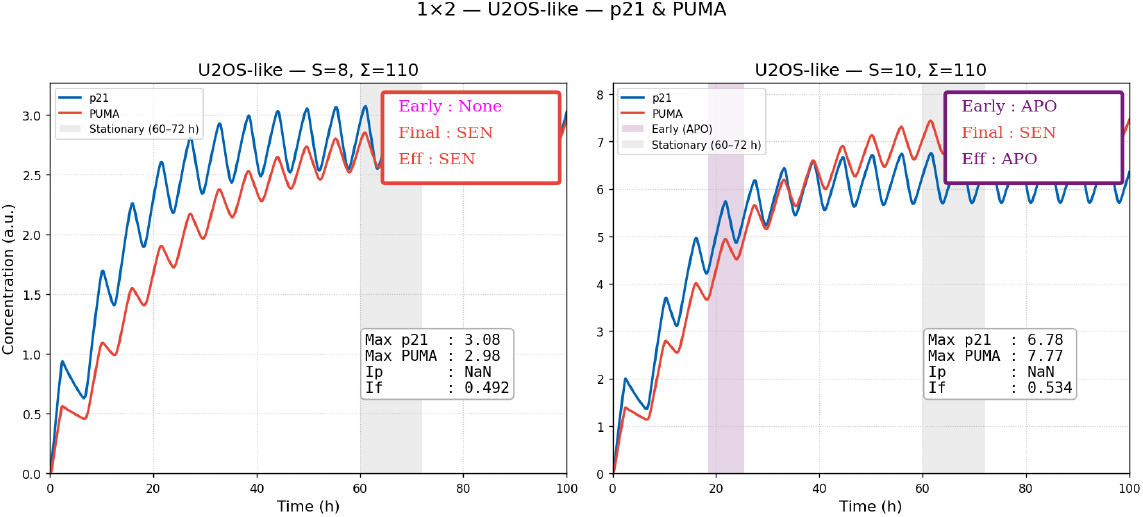
U2OS-like—p21 & PUMA (*S* = 8 and *S* = 10). Gray bands mark the stationary window (60–72 h) and, when present, the colored(purple) band indicates a validated apoptosis early window. **Left (***S* = 8). No early window detected. **Right (***S* = 10). An early PUMA window is crossed.

**Table S4.** U2OS single-cell model parameters used in the simulations (fixed Σ = 110, *S* swept). Decision thresholds are listed elsewhere.

| Parameter | Value | Parameter | Value | Parameter | Value |
| --- | --- | --- | --- | --- | --- |
| $\beta_0$ | 0.0005 | $\beta_{\max}$ | 60 | $K_S$ | 6 |
| $n_\beta$ | 4 | $\delta_p$ | 0.5 | $k_u$ | 30 |
| $K_u$ | 0.15 | $K_0$ | 12 | $n$ | 6 |
| $k_{\text{tr}}$ | 0.07 | $K$ | 12 | $\delta_m$ | 0.4 |
| $k_{\text{tl}}$ | 0.6 | $\delta_M$ | 0.4 | $\xi_1$ | 0.04 |
| $\xi_2$ | 0.40 | $K_1$ | 45 | $K_2$ | 80 |
| $s$ | 4 | $s_1$ | 4 | $\beta_{\text{PUMA}}$ | 0.10 |
| $\varphi_{\text{pu}}$ | 5.6 | $K_a$ | 16.0 | $K_{\text{PUMA}}$ | 9.0 |
| $\delta_{\text{pu}}$ | 0.05 | $\hat{\beta}_{21}$ | 0.06 | $\varphi_{21}$ | 10.50 |
| $K_b$ | 8.0 | $K_{21}$ | 6.8 | $\delta_{21}$ | 0.10 |
| $n_S$ | 2 | $K_{\text{pu}}$ | 3.0 | $S_1$ | 24 |
| $S_2$ | 16 | $n_f$ | 2 | $n_1$ | 12 |
| $\Sigma$ | 110 | $S$ | varied on $[0, 25]$ | | |

The threshold values used for U2OS archetype are listed in the box bellow:

#### Supplementary Box 3.

**U2OS-like** — Decision thresholds (global)

U2OS-decision-critere **Early PUMA (APO):**

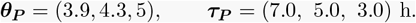

**Stationary gates (mean on** [60, 72] **h):**

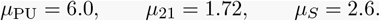

**Index cutoffs (used for** *I*_*f*_ ):

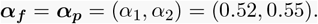

**Early** *p*21 **bands:**

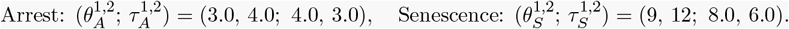

For *S* = 8. No early window is detected (*Early: None*). The 60–72 h averages classify the cell as **senescence** (*Final: SEN* ); the effective decision is **SEN**. There is *no valid early overlap*, hence *I*_*p*_ is undefined. The stationary index is *I*_*f*_ ≃ 0.492. For *S* = 10. A PUMA early window is crossed (*Early: APO*). Stationary averaging yields **arrest** (*Final: ARR*); the effective decision is therefore **APO**. Again there is *no valid early overlap*, so *I*_*p*_ is undefined. The stationary index is *I*_*f*_ ≃ 0.534.

These checkpoints summarize what the figure shows (no *I*_*p*_ at either stress; SEN at *S*=8, early APO at *S*=10) and will be used to introduce the updated U2OS parameter table in Table S4.

Using the decision rule i, we ran population simulations (N=2000 U2OS-like cells) at the literature-supported stress levels (S=0, 8, 10) and compared the simulated fate fractions with published benchmarks; the results are shown in Fig. S11.

**Figure S11.**
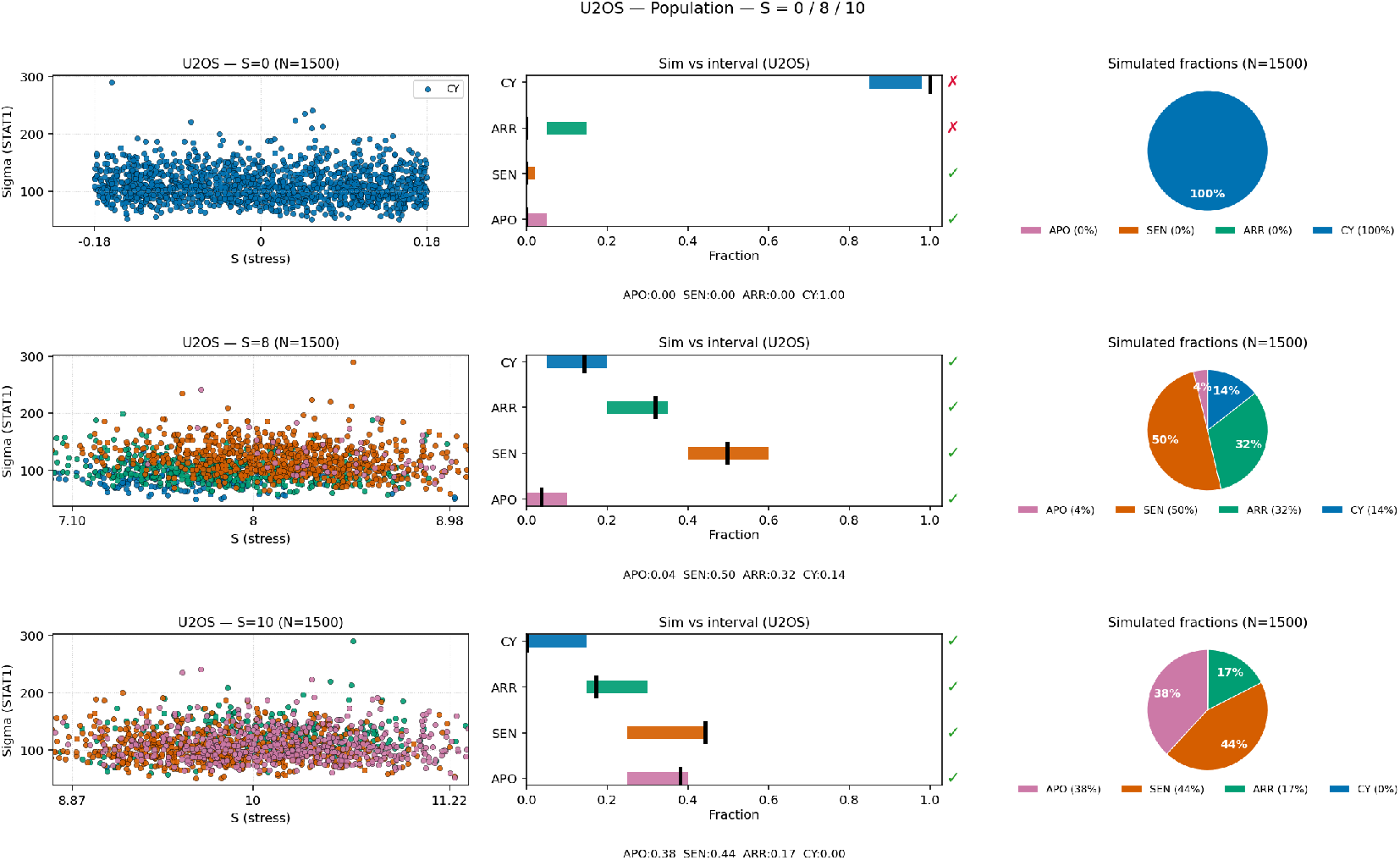
Simulations for the U2OS archetype.

#### Validation of population fractions against literature intervals (U2OS-like)

For each stress level *S* and fate *k* ∈ {CY, ARR, SEN, APO}, the literature specifies an admissible interval [*ℓ*_*S,k*_, *u*_*S,k*_]. From *N* =1500 simulated cells we estimate 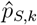, the empirical fraction per fate. On the right-hand panels, colored bars represent the literature intervals, and the black ticks show the simulated 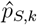; a green 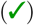 or red 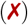 mark indicates whether 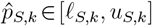.

##### Results summary

Across the three stresses (*S*=0, 8, 10) the validation is globally satisfactory. At *S*=0 the simulation is purely **cycling** (CY= 1.00), matching the high-end of the literature range for viable cycling cells and confirming the absence of stress-induced activation. No senescent or apoptotic response emerges at baseline.

At *S*=8, the distribution becomes mixed, with SEN ≈ 0.50, ARR ≈ 0.31, and a small apoptotic fraction (APO ≈ 0.03). All simulated fractions fall within or touch their literature intervals (**PASS**), indicating a realistic coexistence of transient arrest and senescence. The scatter panel confirms the coexistence of multiple fates within the same (*S*, Σ) range.

At *S*=10, the population shifts further towards SEN and APO (SEN ≈ 0.43, APO ≈ 0.38), with residual ARR ≈ 0.19. All points again lie within the admissible colored bands, giving a full **PASS** classification. This progression (CY → SEN/APO) reproduces the canonical U2OS sequence: cycling cells dominate at low stress, mixed senescence/arrest at moderate stress, and dual senescence–apoptosis responses at high stress.

##### Interpretive remarks

Two aspects are worth noting. (i) The pure-CY outcome at *S*=0 slightly underestimates the minor non-cycling tail allowed by literature, but this likely stems from idealized initial conditions: zero basal *p*53, *p*21, and PUMA, no micro-stress jitter, and narrow STAT1 dispersion suppress early threshold crossings. (ii) At intermediate stress (*S*=8), the simulated fractions sit near the edges of their admissible ranges (ARR at upper bound, SEN at lower), indicating that the model’s decision surfaces are nearly tangent to empirical limits—consistent with the modest normalized gap *G*_*S*_ ≈ 0.2.

Overall, the U2OS-like population exhibits quantitative agreement with the literature intervals across the entire stress range, with only minor edge-touching effects around the intermediate regime where senescence and arrest overlap.

